# Effects of long-term high-temperature stress on reproductive growth and seed development in development in *Brassica napus*

**DOI:** 10.1101/2021.03.11.434971

**Authors:** Kateřina Mácová, Unnikannan Prabhullachandran, Ioannis Spyroglou, Marie Štefková, Aleš Pěnčík, Lenka Endlová, Ondřej Novák, Hélène S. Robert

**Author notes:** Kateřina Mácová, Unnikannan Prabhullachandran, Ioannis Spyroglou, Marie Štefková, Aleš Pěnčík, Lenka Endlová, Ondřej Novák. Hélène S. Robert – corresponding author.

## Abstract

*Brassica napus* is the second most important oilseed crop worldwide. Increasing average temperatures and extreme weather have a severe impact on rapeseed yield. We determined the response of three cultivars to different temperature regimes (21/18 °C, 28/18 °C and 34/18 °C), focusing on the plant appearance, seed yield, seed quality, seed viability, and embryo development. Our microscopic observations identified that embryo development is affected by high temperatures. We noticed an acceleration of its development, in addition to pattern defects. Reduced fertilization rate, increased abortion rate, and preharvest sprouting would be responsible for the low seed yield at the high-temperature regime. Hormone profiling indicates that reduced auxin levels in young seeds may cause the observed embryo pattern defects. Moreover, reduced seed dormancy may result from low ABA and IAA levels in mature seeds. Glucosinolates and oil composition measurements suggest reduced seed quality. These identified cues help understand seed thermomorphogenesis and pave the way to the development of thermoresilient rapeseed plants.

**Highlight:** *Brassica napus* flowering plants’ growth at high temperatures accelerates embryo development, causing a high seed abortion rate and reduced seed quality. Temperature-reduced ABA levels cause premature seed sprouting.

## 1. Introduction

Global climate changes with outbreaks of extreme weather can harm reproductive development and decrease agronomic crops’ overall yield. Severe weather incidents analysed for the period 1964-2007 were shown to profoundly affect crop harvest worldwide due to drought and heat (Lesk *et al*., 2016). This has been reported in rice (Peng *et al.,* 2004; Guo *et al.*, 2019), maize (Liu *et al.*, 2020), wheat (Liu *et al.*, 2016), soybean (Zhao *et al.*, 2017), tomato (Singh *et al.*, 2017), and cotton (Singh *et al.*, 2007). Predicted increase of temperatures in the future will deepen problems with crop yield (Zhao *et al.,* 2017), which encourages scientific efforts in the production of thermotolerant (or stress-tolerant) cultivars. Heat stress (HS) is considered as growing conditions above the critical threshold temperature. For most of the crops, the critical temperature is around 30 °C (Luo, 2011).

Rapeseed is the second most important crop for oil production with high nutritive quality, currently cultivated on 37.58 million ha with an annual production of 75 million tons and average productivity of 1.99 tons/ha (Food and Agriculture Organization of the United Nations, 2020). Total oil production is dependent on several crucial steps leading to plant reproduction and seed production. They comprise the transition to the flowering stage, flower development, ovules and pollen formation, pollination and fertilization processes (ovule viability, pollen germination and pollen tube growth), embryo and endosperm development, and seed development and maturation. The flowering stage is the most temperature-sensitive phase of development (Gan *et al.*, 2004). The critical temperature for rapeseed plants during flowering was suggested to range from 25 °C to 32 °C depending on the studied cultivars (Angadi *et al.*, 2000; Morrison *et al.*, 1989; Polowick *et al.*, 1988). The intraspecific variability of the cultivars in their stress response foresees chances for breeding programmes.

Specific effects of HS on individual generative growth events have been reported in some agronomically important crops. HS reduces pollen viability and fertilization events in pea (Jiang *et al.*, 2019), rice (Das *et al.*, 2014), and chickpea (Devasirvatham *et al.*, 2013). Seed set, seed filling and quality were decreased by HS in wheat (Hays *et al.*, 2007; Talukder *et al.*, 2014), rice (Lin *et al.*, 2010), chickpea (Jumrani and Bhatia, 2014), and sorghum (Impa *et al.*, 2019). Despite many studies reporting damage of pollen grains due to HS in various species, rapeseed pollen manifests a limited HS sensitivity. The average viability of pollen grains decreases from 96.4 % at 25/15 °C to 84 % at 35/25 °C (Chen *et al.*, 2020a). A higher difference in pollen viability (58 % in 35/18 °C compared to 86 % in 23/18 °C) was shown together with reduced pollen germination rates (17.5 % in HS versus 59.2 % in control) (Young *et al.*, 2004). Cross of HS pollen on control pistils resulted in a markedly decreased number of seeds compared to control. Similar observations but to a lesser extent were made for the reciprocal cross (Young *et al.*, 2004). HS’s effects on female gametophyte and early embryo development, which could explain the remaining variability in the decreased yield production, have yet to be elucidated. Seed weight and seed number per pod were reduced in cultivation above 29.5 °C of canola cultivars (Morrison and Steward, 2002). Reduced pod number and seed yield were caused by 32/22 °C and 35/25 °C day/night temperatures in a 7-day treatment during the week before and after flowering initiation of 12 different cultivars (Chen *et al.*, 2020a). High variability among cultivars was observed for pollen viability and germination in the temperature range 5-35 °C with a significant decrease above 30 °C (Singh *et al.*, 2008). Total yield, seed number and seed weight were also reduced by a two-week 31/14 °C treatment (Koscielny *et al.*, 2018). High-temperature treatments induce changes in seed oil content and photosynthesis activity. It suggested that the damage to PSII and inhibition of a fatty acid biosynthesis pathway controlled by the transcription factor BnWRI1 might be the major causes of the decreased oil content of *B. napus* seeds that develop under HS (Huang *et al.*, 2019).

The studies mentioned above are focused on an HS lasting from hours to several days. The impact of long-term elevated temperatures above critical temperature threshold during the whole reproductive phase was performed on *B. napus* cultivar (N99-508) at 29 °C (Elferjani and Soolanayakanahally, 2018), with a similar outcome as short treatment, e.g., altered photosynthesis, reduced seed yield and seed oil content (Huang *et al.*, 2019; Koscielny *et al.*, 2018). In the cultivar Aviso, no significant difference in yield was observed at 33 °C. Nevertheless, HS altered seed quality and induced pre-harvesting seed sprouting, correlating with decreased abscisic acid (ABA) levels in mature seeds (Brunel-Muguet *et al.*, 2015). In response to stress, plants change their growth by altering homeostasis, transport and signalling of phytohormones. Elucidation of crosstalk of phytohormones and regulation of morphogenesis under HS (thermomorphogenesis) is still in progress in many different crops and tissues (Torres *et al.*, 2017; Stavang *et al.*, 2009). ABA is responsible for stress tolerance and acts as a regulator for responding to various stress, including HS (reviewed in Tuteja, 2007; Vishwakarma *et al.*, 2017). ABA controls various processes in plants, e.g., stomatal closure and water transport (Parent *et al*., 2009; Christmann *et al.*, 2007), maintenance of seed dormancy and control of seed germination by crosstalk with gibberellic acid (Seo *et al.*, 2006). ABA-mediated heat tolerance may function by regulating ABA-responsive genes containing ABA-responsive elements in their promoter regions, which subsequently activate heat shock transcription factors (HSFs) and heat shock proteins (HSPs). ABA increases the production of HSPs (and H_2_O_2_) in maize (Hu *et al.*, 2010), wheat (Hu *et al.*, 2018), rice (Zhang *et al.*, 2014), and cucumber (Li *et al.*, 2014). Moreover, two ABA-responsive genes (*FaAREB3* and *FaDREB2A*) may act as regulators of *FaHSFA2c* and the downstream *FaHSPs* in tall fescue (*Festuca arundinacea*) (Wang *et al.*, 2017). Some HS-related genes, including HSP, HSF and other seed-related genes of rapeseed activated by HS, were identified (Yu *et al.*, 2014; Young *et al.*, 2004).

Stress response promotes secondary metabolites production. Indole glucosinolates are specific secondary metabolites of the *Brassicaceae* family derived from tryptophan, connected to auxin biosynthesis, and involved in stress response (Eom *et al.*, 2018; Salehin *et al.*, 2019) and secondary seed dormancy (Liu *et al.*, 2019). It has been shown in Arabidopsis that glucosinolates may trigger thermotolerance by inducing HSP and H_2_O_2_ (Ludwig-Müller *et al.*, 2000; Hara *et al.*, 2013). However, its detailed regulation is still unknown and requires further investigation in specific developmental stages.

Environmental signals, phytohormones, and abiotic stresses are integrated with the control of circadian rhythm. Genes related to circadian clocks encode transcription factors arranged in several feedback loops to synchronize plant physiology with the daily changing environment (Sanchez *et al.*, 2011). Circadian gene expression was altered by both cold treatment (Bieniawska *et al.*, 2008) and heat treatment in Arabidopsis and other crops (Blair *et al.*, 2019; Li *et al.*, 2019). Moreover, alternative splicing plays a role in development, stress response, and circadian clocks regulation (Sugliani *et al.*, 2010; Fouquet *et al.*, 2011; James *et al.*, 2012; Filichkin *et al.*, 2015).

This study investigated the effects of two high-temperature regimes on three *B. napus* cultivars focusing on reproductive growth, especially changes in maternal tissue, early seed and embryo development, and seed yield and quality. We measured yield traits and supplemented these macroscopic data with a detailed analysis of seed and embryo development to examine embryo defects and analysis of seed germination and seedlings. We performed hormonal and seed composition profiling and identified development- and hormone-related deregulated genes, which could explain the plant HS response in specific development steps and outline possible targets for yield improvement.

## 2. Materials and methods

### 2.1 Plant materials and growing conditions

Three *Brassica napus* cultivars (Westar, Topas and DH12075) were tested in three chambers with different temperature conditions, ten plants per cultivar in four biological replicates, in greenhouses of CEITEC MU Plant Sciences Core Facility. Seeds were sterilized by 20 % bleach, washed twice in sterile distilled water, vernalized at 4 °C for 24 hours, germinated on plates containing MS medium for five days (21 °C, long day, 150 uE) and transferred to soil in 0.7 L pots. After two weeks in phytotron (21 °C, long day, 150 uE), the plants were fertilized with KRISTALON™ Start (N-P-K (19-6-20) + 3 % Mg + 7.5 % S) and one week later transferred to 1.5 L pots. With the first visible flowering buds, the plants were randomly transferred to greenhouse chambers with a long day regime, LED lights with light intensity 150 uE, 35-45 % humidity, 18 °C during the night. Temperature condition for control (CT), mid-(MT) and high-temperature (HT) chambers were set to 21 °C, 28 °C and 34 °C, respectively, with ramping of the temperature up and down by 4 °C per hour (Supplementary Fig. S1 at *JXB* online). Plants were well watered to avoid any effect associated with drought stress. Plants were once fertilized with KRISTALON™ Fruit and Flower (N-P-K (15-5-30) + 3 % Mg + 5 % S) at the flowering start and kept in the respective chamber till seed harvest. The experiment was performed during wintertime to reduce external influences to a minimum.

### 2.2 Plant phenotyping measurements

The number of leaves (NL) was counted at the beginning of flowering as a control trait. The length of the main flowering stem (LMS, cm), the duration of flowering time (FT, days), the number of primary branches on the main stem (NB), the number of flowers on the main stem (NF) and the number of ovules per pistil were evaluated during each experiment (four replicates, ten plants per cultivar). LMS is the difference in length of the main stem between the beginning and the end of the flowering period. For silique length (SL, cm), 30 flowers were hand-pollinated for each cultivar, the SL was measured daily for 11 days, and the number of seeds (NS) was counted for correlation analysis.

Pollen viability was tested by both acetocarmine and Alexander staining. Alexander staining solution was prepared according to Alexander (1969). Pollen grains were tapped on a microscopic slide with 10 μL of Alexander solution, incubated at 50 °C for 24 hours and counted under the light microscope Zeiss Axioscope.A1. For acetocarmine solution, 2 % carmine solution was made in 95 % glacial acetic acid (Marutani *et al*., 1993). Pollen grains were stained for 10 minutes and evaluated using the same microscope.

Silique length was measured for five siliques on six plants for each biological replicate each day till 11 days after pollination (DAP). The final length and number of seeds per silique were recorded at harvest. Siliques with a length < 1 cm were excluded (e.g., no seeds). For examining the temperature effect on silique growth after pollination, the Relative Growth Rate (RGR) from the exponential growth model was used. RGR is estimated by the coefficient (r) from the exponential function: Y=C*e^rt^, where Y is the variable modelled and C, a constant term. When r = 0.1, the silique grows by an average of 10 % every time point t.

### 2.3 Seed phenotyping

Seeds from five hand-pollinated flowers on ten plants were evaluated for the number of seeds and different phenotypes of mature seeds after harvest. Moreover, seeds from two plants were collected at 3, 4, 5, 6, 7, and 8 DAP and cleared in a chloral hydrate solution (chloral hydrate, Sigma C8383/Glycerol/Water, 8/1/3 w/v/v) for embryo phenotyping. Observations were made using a light microscope Zeiss Axioscope.A1 equipped with DIC optics and ZEN blue software for picture analysis at CEITEC MU Cell Imaging Core Facility. These siliques were also used for the number of ovules (NO), fertilization efficiency and abortion rate assessment. Premature seed germination (sprouting, PHS) was evaluated at 26 DAP. After harvest, seeds were stored in paper bags at room temperature. The 100-seed weight and germination rate with seedling viability assessment were analysed four months after harvest.

### 2.4 Data analysis

The results (multiple pairwise comparisons between different conditions-temperatures) were analysed using generalized linear mixed models (Poisson or negative binomial mixed model for count data and linear mixed model for continuous data), and analysis of variance (ANOVA) followed by Tukey’s posthoc test in R software (R Core Team, 2014) in RStudio (RStudio Team, 2019). Mixed models were used, so the randomness between the different batches is considered (McCulloch and Neuhaus, 2005). The level of statistical significance was set at p ≤ 0.05 for all tests. Packages “glmmTMB” and “emmeans” were used for the fitting of mixed models and implementing pairwise comparisons, respectively (R Core Team, 2014; Brooks *et al*., 2017; RStudio Team, 2019; Russell, 2020).

### 2.5 Seed metabolites content

Harvested mature seeds were analysed for their oil composition and glucosinolates content. Three technical and two biological replicates from CT, MT, and HT regimes were analysed except for Westar and DH12075 from HT, where only one biological replicate was used due to low seed production. Sample measurements were performed by a spectrophotometer FT-NIR Antaris II (Thermo Fisher Scientific Inc., USA) on the integration sphere in reflectance mode in a spectral range of 10 000 - 4 000 cm^−1^ using OMNIC for Antaris software. Whole seeds were measured in rotary circular cuvettes with quartz bottom permeable for NIR radiation. The resulting spectrum of each sample was obtained as an average of 64 scans with a resolution of 2 cm^−1^. Calibration models for quantitative analysis of the oil, main fatty acids (palmitic, stearic, oleic, linoleic, linolenic, erucic), and glucosinolates content were developed using Partial Least Squares algorithm in Thermo Scientific TQ Analyst software. To construct FT-NIR calibration models, data measured by routine laboratory reference methods were used. Determination of oil content was performed by extraction method according to ČSN EN ISO 659 (2009), which defines the weight determination of oil content after extraction with petroleum ether, distillation of the solvent and drying the extracted fat. The dry matter content was determined by a gravimetric method, according to ČSN EN ISO 6652 (2001) after 4 hours of drying at 103 °C. To determine the GSL content, the HPLC / UV-VIS method, according to ČSN EN ISO 9167-13 (1998), was used, which defines the GSL determination in the form of desulfoglucosinolates. The representation of individual fatty acids in the form of fatty acid methyl esters was detected by GC/FID according to the internal methodology of the Research Institute of Oilseed Crops. All methods used are validated and routinely used. Data of total oil are presented as % content in dry matter and in tissue at 8 % humidity. The content of each fatty acid is shown as % of the total oil content. The number represents the mean value ± S.D. Statistical evaluation for significant difference in heat treatments was performed by a paired Student’s t-test (*, **, and *** correspond to P-values of 0.05 > p > 0.01, 0.01 > p > 0.001, and p < 0.001, respectively).

### 2.6 Hormonal measurements

IAA, its metabolites, and ABA were determined following the methods described by Pěnčík *et al*. (2018). Samples containing 10 mg of frozen homogenized tissue were extracted with 1 mL of 50 mM phosphate buffer (pH 7.0) containing 0.1 % sodium diethyldithiocarbamate and a mixture of stable isotope-labelled internal standards. One portion of the extract (200 μL) was acidified with HCl to pH 2.7 and purified by in-tip micro solid phase extraction (in-tip μSPE). Another 200 μL of the extract were purified directly without acidification to determine IAOx. The last 200 μL portion was derivatized by cysteamine, acidified with HCl to pH 2.7 and purified using in-tip μSPE in order to determine IPyA. After evaporation under reduced pressure, samples were analysed using HPLC system 1260 Infinity II (Agilent Technologies, USA) equipped with Kinetex C18 column (50 mm x 2.1 mm, 1.7 μm; Phenomenex) and linked to 6495 Triple Quad detector (Agilent Technologies, USA). Data are presented as pmol for 1 g of fresh tissue. The number represents the mean value ± S.D. Statistical evaluation for significant difference in heat treatments was performed by a paired Student’s t-test (*, **, and *** correspond to P-values of 0.05 > p > 0.01, 0.01 > p > 0.001, and p < 0.001, respectively).

### 2.7 RNA extraction and RT-qPCR

Tissue (50 to 100 mg per replicate) collected from CT and HT regimes was directly frozen and stored in a −80 °C freezer. For isolation, samples were ground into a fine powder using mortar or ceramic beads added to 2 mL Eppendorf tubes. The total RNA from leaves, pistils and young seeds (7 DAP from CT and 5 DAP from HT) were isolated following a protocol for Trizol (Invitrogen). RNA from 26 DAP seeds was isolated using a NucleoSpin RNA Plant and Fungi kit (Macherey-Nagel) according to the manufacturer’s protocol. All samples were treated with a rDNase, RNase-free (Macherey-Nagel). M-MLV Reverse Transcriptase (Promega) was used for reverse transcription on two μg of DNA-free RNA.

Expression of genes involved in auxin and ABA biosynthesis, degradation and signalling, and genes related to mRNA splicing pathway and regulators of flowering and thermogenesis were quantified. Gene sequences for primers design were available in the Genebank database (LOC number listed in Supplementary Table S1). The selection of the candidate reference genes for normalization was made using the combination of the algorithms provided by Normfinder (Andersen *et al.*, 2004), geNorm (Vandesompele *et al.*, 2002) and BestKeeper (Pfaffl *et al.*, 2004). The geometric mean of the selected genes was then used as a normalization factor for calculating the log2-fold change. The selected reference genes were *BnaACT7* (leaves, pistils, and 26 DAP seeds), *BnaeIF5A* (leaves, pistils, young seeds, and 26 DAP seeds) and *BnaTMA7* (pistils and young seeds). All primers sequences used in this study are listed in Supplementary Table S1. Gene transcript abundance was quantified by RT-qPCR using FastStart Essential DNA Green Master (Roche) on a Lightcycler 96 (Roche). The experiment was performed with three biological replicates, each with three technical replicates. Analysis of log2-fold changes was performed using the 2^ΔΔCt^ method. An expression change is estimated as significant with a log-fold change of ± 0.8. The qPCR reactions were performed according to MIQE Guidelines (Bustin *et al.,* 2009).

## 3. Results

### 3.1 Effect of high-temperature treatment on overall plant growth

Three *Brassica napus* cultivars were evaluated for main growth characteristics in the three temperature regimes. At the beginning of the experiment, NL (number of leaves) was evaluated with no significant difference among the temperatures demonstrating a comparable vegetative growth of all the plants randomly distributed to the different greenhouse chambers (Supplementary Fig. S2A). For all the cultivars, significantly higher LMS (length of the main stem) and NF (number of flowers) were detected in HT conditions (Figs 1A, 1D-F, Supplementary Fig. S2B). The flowering stem’s growth speed is more accelerated than the production of flowers by the floral meristem on the main stem, resulting in a decreasing trend in NF/LMS ratio (Fig. 1B). A strong correlation was detected in CT (control growth temperatures, 21/18 °C) between LMS and NF with a comparable level among all three cultivars (> 0.77). In MT (mid growth temperature, 28/18 °C), the strong correlation was maintained in Topas (0.90) and Westar (0.74) but dropped to 0.43 in DH12075. In HT (high growth temperature, 34/18 °C), on the contrary, this correlation dropped for Topas (0.44) to almost no correlation for Westar (0.22) and increase to 0.77 for DH12075 (Supplementary Table S2).

**Fig. 1.**
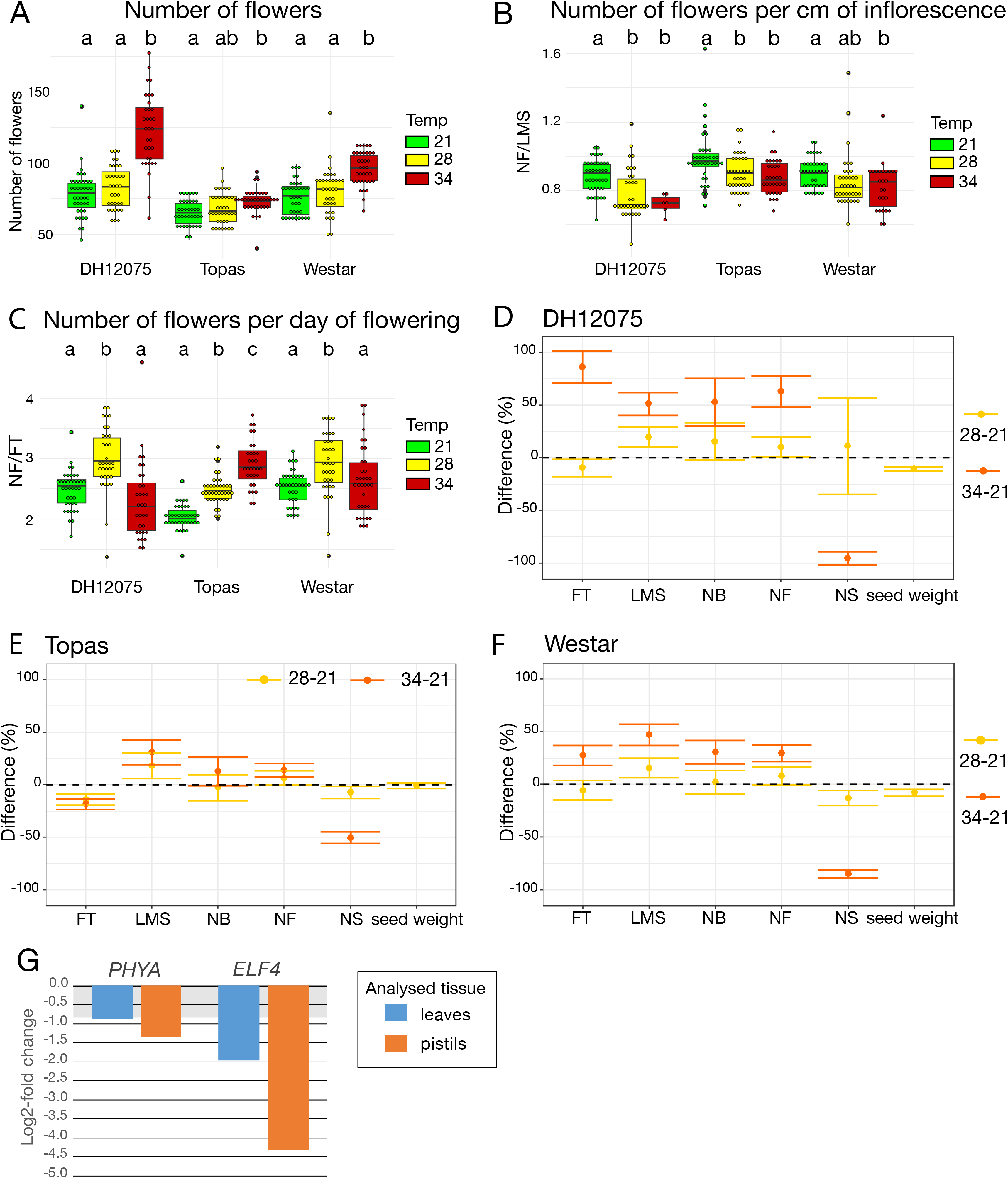
Effects of high temperatures on flowering traits and seed production (A-C) The number of flowers (NF) (A), number of flowers per cm of inflorescence (NF/LMS) (B), and number of flowers per day (NF/FT) (C) were quantified in DH12075, Topas and Westar cultivars at CT (21/18 °C, green), MT (28/18 °C, yellow) and HT (34/18 °C, red). The number of flowers is presented as boxplots. The box represents the interquartile range, and the line inside the box represents the median. The Pearson correlation coefficient between LSM, FT, and NF, is shown in Supplementary Table S2. Boxes with the same letters (a, b, c) within each cultivar do no differ significantly (p < 0.05). (D-F) Graphs are displaying the differences between CT and MT (yellow), and CT and HT (orange) in flowering time (FT, days), length of the main stem (LMS, cm), number of branches (NB), number of flowers (NF), number of seeds (NS) and seed weight, in DH12075 (D), Topas (E) and Westar (F). Dots represent the mean values of the differences and error bars, the 95 % confidence interval. Data on FT, LMS, and NB are presented in Supplementary Fig. S1. Data on NS and seed weight are presented in Fig. 3. (G) Expression analysis by RT-qPCR of *BnaPHYA* and *BnaELF4* in Westar leaves (blue) and pistils (orange). The graphs are displaying the log2-fold changes in expression between CT and HT. The grey zone covers the [−0.8 to 0] log2-fold changes, considered a non-significant change. Primers and LOC information are presented n Supplemental Table S1.

Differential growth imposed by high temperatures was also reflected in FT (flowering time). In MT, Westar and Topas reacted with shortening their flowering time (Figs 1E-F, Supplementary Fig. S2C). Topas kept a similar FT in HT and MT due to problems with the floral meristem, restricting growth. DH12075 and Westar displayed a significantly longer FT in HT (Figs 1D-F, Supplementary Fig. S2C). Therefore, these two cultivars developed more flowers per day in MT but not HT (Fig. 1C). On the other hand, due to a shorter FT, Topas had a significantly higher NF/day in MT and HT (Fig. 1C). In general, a strong correlation was found between FT and NF in CT and MT, with a decreasing tendency to a low correlation in HT (Supplementary Table S2). Moreover, all the cultivars developed more branches in HT treatment (Supplementary Fig. S2D).

Longer main stems, prolonged duration of flowering and more branches in DH12075 and Westar cultivars (Figs 1D, 1F) implicate changes in hormonal levels, maintenance of growth (reduced senescence/dormancy) and decreased apical dominance in HT growth conditions. On the other hand, Topas developed more flowers per day due to a floral meristem arrest at HT. However, looking at the other measured parameters, Topas appeared less negatively affected by high temperatures on the whole plant level than the other cultivars (Fig. 1E).

The relative expression of two genes related to environmental conditions sensing and circadian clock regulation was assessed. Both *PHYTOCHROME A* (*PHYA*), encoding a photoreceptor, and *EARLY FLOWERING 4* (*ELF4*) control induction of flowering and regulate circadian clock loops (McWatters *et al.*, 2007; Seaton *et al.*, 2018). Moreover, *PHYA* and *ELF4* are temperature-responsive (Song *et al.*, 2017; Song *et al.*, 2018; Chen *et al.*, 2020b). In accordance, we found both *PHYA* and *ELF4* to be significantly downregulated in HT in leaves and pistils (Fig. 1G). A lower amount of *PHYA* and *ELF4* may slow down the regulation of the circadian clock. However, their specific effects on the FT, NB, or LMS have to be further evaluated, even though circadian clocks’ general effect on plant architecture has been studied (Rubin *et al.*, 2018).

### 3.2 Pollination and embryo development

The success of reproduction relies on the regulated development of ovules within the pistil and pollen grains inside the anthers. It is also determined by the proper timing of pollination and fertilization events for subsequent seed development.

The NO (number of ovules per pistil) was comparable in all the temperature regimes (Table 1, Supplementary Fig. 3A). For Westar, we found a minimal but significant difference between CT and HT, which may be caused by the higher variability of this trait in Westar cultivar, but its biological relevance is relatively small.

**Table 1.** Number of ovules per pistil and outcome after fertilization

| Conditions |  | NO* | Seeds (%) | Non-fertilized (%) | Aborted (%) |
| --- | --- | --- | --- | --- | --- |
| DH12075 | CT | 24.78 – 27.66 | 90.81 | 8.05 | 1.14 |
|  | MT | 24.29 – 27.39 | 78.06 | 19.45 | 2.50 |
|  | HT | 26.05 – 28.79 | 47.27 | 51.81 | 0.93 |
| Topas | CT | 21.33 – 24.05 | 89.25 | 7.41 | 3.34 |
|  | MT | 21.98 – 24.53 | 79.09 | 14.30 | 6.61 |
|  | HT | 22.65 – 25.03 | 38.58 | 46.12 | 15.30 |
| Westar | CT | 26.31 – 33.45 | 92.13 | 6.84 | 1.03 |
|  | MT | 24.78 – 31.19 | 69.59 | 20.95 | 9.46 |
|  | HT | 23.57 – 29.96 | 32.55 | 55.05 | 12.40 |

\* 95% confidence interval

Pollen viability decreased by less than 5 % by temperature treatment, which was not considered a significant effect (Supplementary Fig. S4). Fertilization and abortion rates were monitored from 5 to 8 DAP. In all three cultivars, the percentage of non-fertilized ovules increases with increasing temperature resulting in a reduced number of developing seeds. Moreover, Topas and Westar cultivars also exhibited an increasing percentage of aborted seeds. On the contrary, the abortion rate in DH12075 decreased in HT compared to CT and MT (Table 1).

Seeds were investigated to monitor embryo development from 3 DAP to 8 DAP (Table 2). At 3 DAP, all the cultivars exhibited embryos in the zygote or one-cell stage. The embryo development is accelerated with increasing temperatures, leading to two development stages difference at 6 DAP (Table 2, Supplemental Figs 3B-J). Faster embryonic development may affect the synchronous development of the seed coat and the endosperm and eventually lead to a higher abortion rate in later stages (Table 1). Less than 1 % of defective embryos were observed in CT for all cultivars. This number significantly increased for DH12075 (6 %) and Westar (10 %) in MT, and for all the cultivars in HT (DH12075, 25.45 %; Topas, 16.2 %, and Westar, 39.58 %) (Fig. 2A).

**Table 2.** Most abundant embryo stages per DAP (>20% of the embryos per silique at the given DAP)

| Conditions |  | Embryonic stage |  |  |  |  |  |  |
| --- | --- | --- | --- | --- | --- | --- | --- | --- |
|  |  | Zygote | 1-cell | 2-cell | 8-cell | EG | MG | LG |
| DH12075 | CT | 3 | 3-4 | 5-6 | 6-7 | 7-8 | >8 | >8 |
|  | MT | 3 | 3 | 4 | 4-5 | 5-6 | 6 | 7-8 |
|  | HT | 3 | 3 | 4 | 4-5 | 5 | 6 | 7 |
| Topas | CT | 3 | 3-4 | 4-5 | 5-6 | 6-7 | 7-8 | 8 |
|  | MT | - | 3 | 4 | 4-5 | 5 | 6 | 6-7 |
|  | HT | 3 | 3 | 3-4 | 4 | 5 | 5-6 | 6 |
| Westar | CT | 3 | 3-4 | 5 | 5-6 | 7 | 8 | 8 |
|  | MT | 3 | 3-4 | 4 | 5 | 6 | 6 | 7-8 |
|  | HT | - | 3 | 3 | 4 | 5 | 5 | 6 |

6 DAP is highlighted to illustrate the shift in the embryo development timing.

**Fig. 2.**
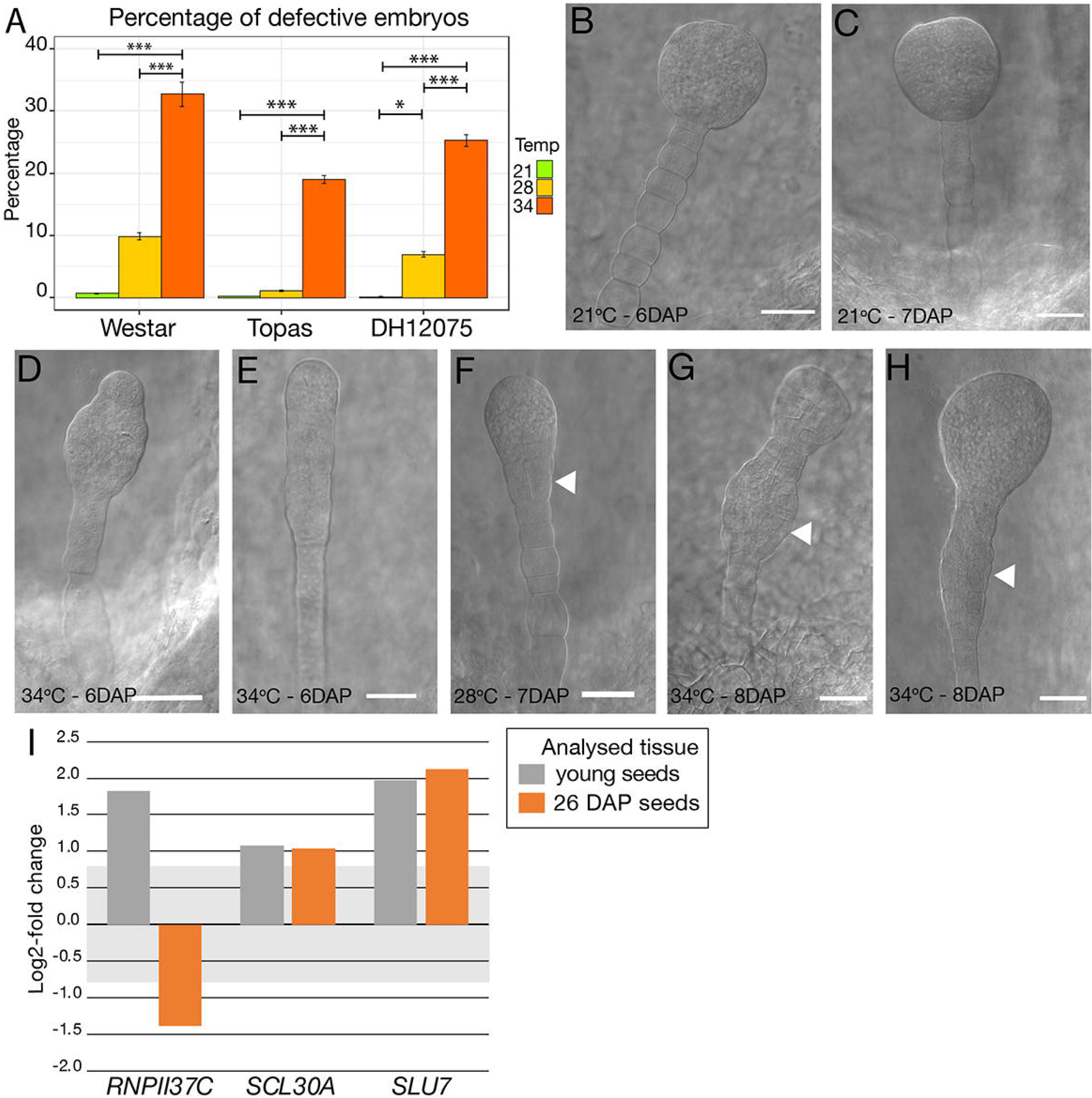
Effects of high temperatures on embryo development (A) Graph displaying the percentage of defective embryos in young seeds of Westar, Topas, and DH12075 plants grown at CT (21/18 °C, green), MT (28/18 °C, yellow) and HT (34/18 °C, orange). Error bars represent the 95 % confidence interval. Asterisks indicate statistically significant difference in MT and HT in an analysis of variance (ANOVA) followed by Tukey’s posthoc test (* and *** correspond to P-values of 0.05 > p > 0.01 and p < 0.001, respectively). (B-C) Embryos at 6 DAP (B) and 7 DAP (C) from plants grown at CT (21/18 °C). (D-H) Range of defective embryos observed in *B. napus* plants grown at MT (28/18 °C) and HT (34/18 °C) between 6 and 8 DAP. The white arrowheads (F-H) point to cell proliferation in the suspensor that may lead to a secondary embryo (G). Scale bars represent 50 μm (B-G). Uncropped pictures are presented in Supplementary Fig. S5. (I, J) Expression analysis by RT-qPCR of *BnaRNPII37C*, *BnaSCL30A* and *BnaSLU* in Westar young seeds (grey) and 26 DAP seeds (orange). The graphs are displaying the log2-fold changes in expression between CT and HT. The grey zone covers the [−0.8 to 0.8] log2-fold changes, considered a non-significant change. Primers and LOC information are presented n Supplemental Table S1.

The defective embryos exhibited several altered cell division patterns in the basal embryonic part and cell proliferation in the suspensor (Figs 2B-H, Supplementary Fig. S5). Moreover, we observed secondary embryos emerging from the proembryo or the suspensor (Fig. 2G), leading probably to twin embryo development. Aberrant or misregulated cell divisions in the apical embryo domain cause incomplete cotyledon development (Fig. 2H). The appearance of aberrant embryos denotes HT’s influence on auxin homeostasis, transport or signalling, as such phenotypes were observed in Arabidopsis mutants disrupted in those auxin-related processes (Robert *et al*., 2013; Prigge *et al*., 2020; Radoeva *et al*., 2019).

These data demonstrated that the decrease in rapeseed yield due to warm growth temperatures is mostly caused by decreased fertilization rate and defective embryo development.

### 3.3 Seed phenotyping

#### 3.3.1 Seed yield

Silique length (SL) is believed to correlate with the number of seeds they contain (Bac-Molenaar *et al.*, 2015). We have noticed that about 50 % of the ovules are not fertilized and that only 32-47 % of the ovules will develop into seeds (Table 1). Therefore, siliques were assessed to their growth rate for 11 days from hand pollination. Siliques from all three cultivars grew on average by 22-23 % daily at CT during the 11 days (Supplementary Fig. S6). This growth rate significantly decreased when growth temperature increased to reach a daily growth rate of 18-20 % and 10-15 % at MT and HT, respectively, for all cultivars. The growth rate at HT is, therefore, twice slower than at CT. There is a strong correlation (0.92-0.96 and 0.92-0.99) between the reduced seed number per silique and the reduced growth rate of the siliques at MT and HT, respectively, compared to CT (0.63-0.83) (Supplementary Table S3).

The number of seeds per silique and seed aspect were examined after harvest, as some seeds did not correctly develop (Table 1, Fig. 2). Three categories of seeds: fully filled (Fig. 3A), partially filled/shrunken (Figs 3B-D), and germinated (Figs 3E-F), were considered viable seeds. High temperatures severely affected the viable seed yield, with a significant decrease at MT, and a critical one at HT, resulting in almost no viable harvested seeds (Fig. 3G). The Topas cultivar has the highest number of germinated seeds before harvest (preharvest sprouting, PHS). It was evident that part of those seeds germinated even before maturation (Figs 3E,F, Figs 4A,B, Table 3).

**Fig. 3.**
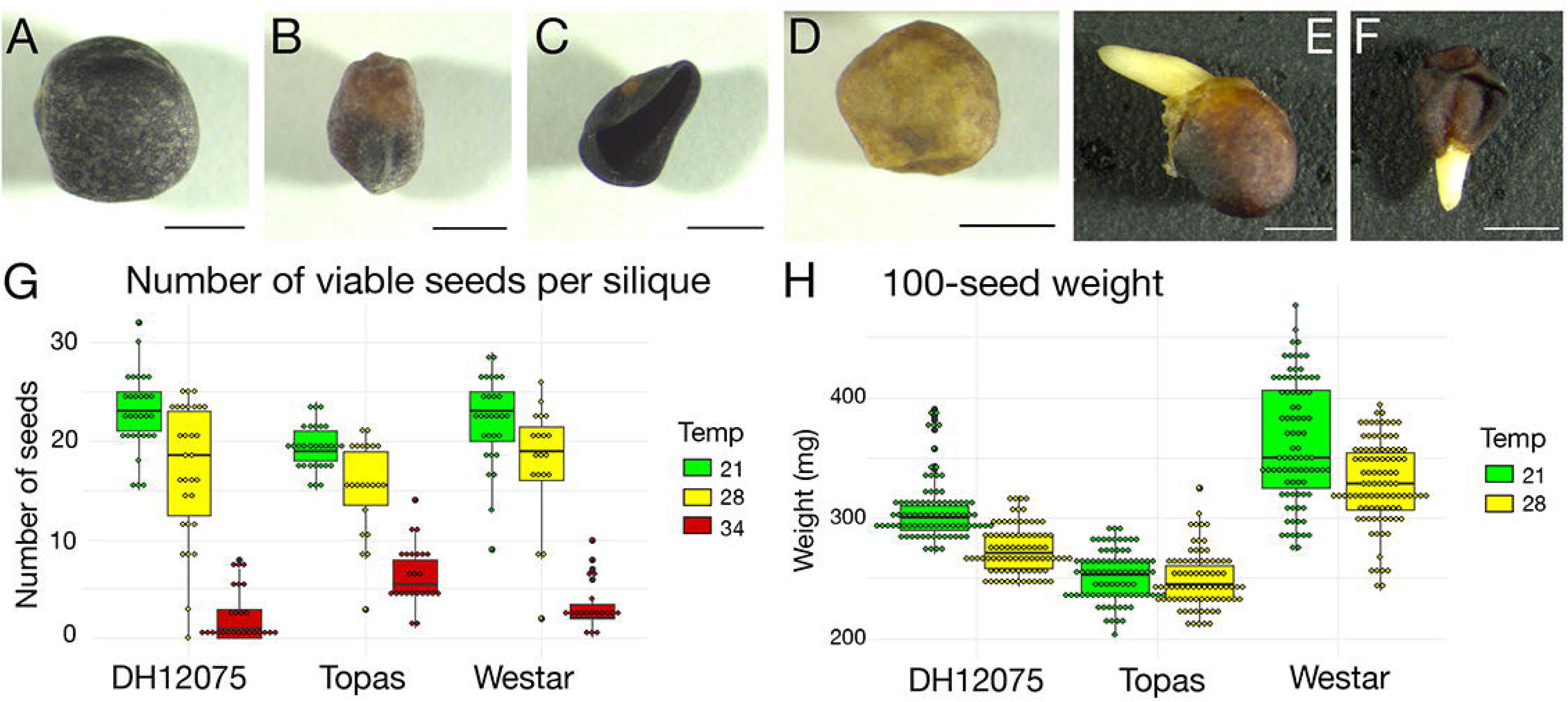
Effects of high temperatures on seed development (A-F) Range of phenotypes observed in mature seeds. We observed normal fully filled seed (A), partially filled seed (B), flatten shrunken seed (C), partially filled yellow seed (D), sprouting fully filled seed (E), sprouting flatten shrunken seed (F). Scale bars represent 1 mm. Graph displaying the number of viable seeds per silique in DH12075, Topas and Westar cultivars at CT (21/18 °C, green), MT (28/18 °C, yellow) and HT (34/18 °C, red). (H) Graph displaying 100-seed weight in DH12075, Topas and Westar cultivars at CT (21/18 oC, green) and MT (28/18 °C, yellow). The data (G, H) are presented in boxplots. The box represents the interquartile range, and the black line inside the box represents the median.

**Table 3.** Comparison of pre-harvest germination of mature and dry seeds (percentage of total seeds for one plant)

|  | 21 °C |  | 34 °C |  |
| --- | --- | --- | --- | --- |
|  | 26 DAP | After harvest | 26 DAP* | After harvest |
| DH12075 | 0 | 0.53 | 2.8 – 13.2 | 11.9 |
| Topas | 0 | 0.7 | 30.4 – 73.4 | 40.8 |
| Westar | 0.45 | 0.22 | 3.8 – 7.9 | 4.1 |

\*95% confidence interval

**Fig. 4.**
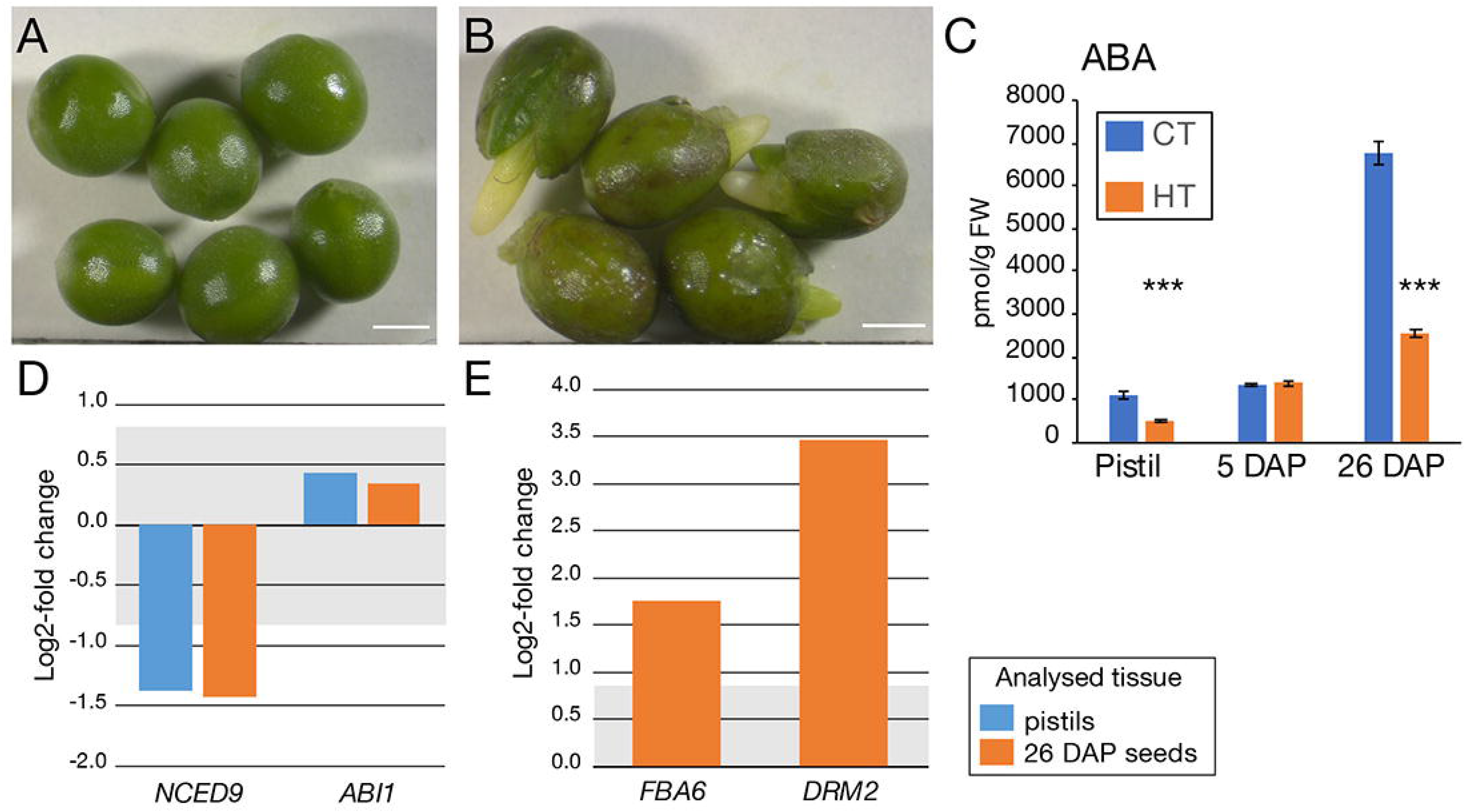
Preharvest sprouting seed phenotypes may be linked to a decrease in ABA levels (A, B) Topas 26 DAP seeds grown at CT (21/18 °C) (A) and HT (34/18 °C). Notice the sprouting of the seeds in B, with the emergence of either root or cotyledons. Scale bars represent 1 mm. (C) Graph showing the ABA levels (in pmol/g FW) in Westar pistils, 5 DAP seeds and 26 DAP seeds grown at CT and HT. Shown is the average ± S.D. Asterisks indicate statistically significant difference in HT in a paired Student’s t-test (t-test; *** corresponds to P-values of p < 0.001). Source data are presented in Supplementary Table S6. (D, E) Expression analysis by RT-qPCR of *BnaNCED9*, *BnaABI1*, *BnaFBA6* and *BnaDRM2* in Westar pistils (blue, D, E) and 26 DAP seeds (orange, E). The graphs are displaying the log2-fold changes in expression between CT and HT. The grey zone covers the [−0.8 to 0.8] log2-fold changes, considered as non-significant changes. Primers and LOC information are presented n Supplemental Table S1.

Fully and partially filled seeds were investigated for seed weight characteristics. In DH12075 and Westar, the 100-seed weight decreased significantly in MT conditions (Fig. 3H). A decrease in weight was not identified in Topas. With the highest amount of seeds per silique and stable seed weight, Topas could be considered the best yield provider under our suboptimal growth conditions.

#### 3.3.2 Preharvest sprouting (premature seed germination)

Based on the mature seed phenotyping analysis, a high amount of Topas seeds displayed a sprouting phenotype, which appeared before seed maturation. In these seeds, either the embryo emerged from the seed coat or the seed coat ruptured due to the embryo pressure from 20 to 35 DAP. PHS rarely occurred in CT (less than 1 % for all cultivars), but its rate is significantly higher in HT (30.4-73.4 % in Topas) (Table 3). To a lesser extent, it was also observed in Westar (3.8-7.9 %) and DH12075 (2.8-13.2 %) in HT (Table 3). The occurrence of PHS corresponds with the percentage of germinated dry seeds after harvest.

#### 3.3.3 Seed germination and seedling viability

Fully and shrunken filled seeds were investigated for germination. Seedling viability and appearance were monitored for seeds collected in CT and MT. In general, most of the seeds (> 80 %) germinated one day after vernalization, regardless of the temperature the seeds developed, reaching > 94 % of germination rate after the 4^th^ day. Within this range, a reduction in germination was notable for DH12075 (2 %) and Westar (4 %) seeds (Fig. 5A).

**Fig. 5.**
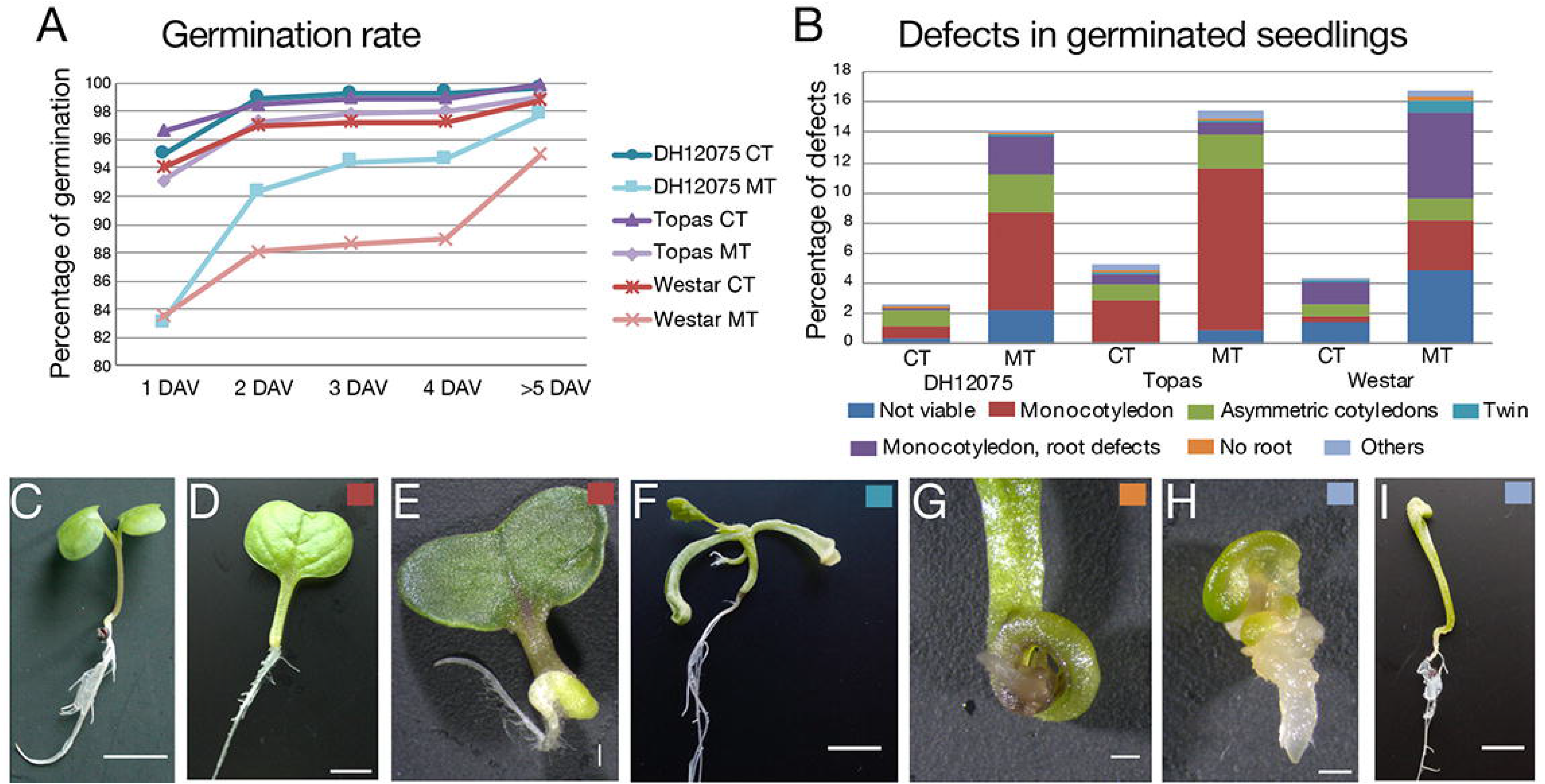
Seed development at high temperatures affects seedling viability. (A) Graph displaying the cumulative percentage of seed germination at 1 Day after Vernalization (DAV), 2 DAV, 3 DAV, 4 DAV and later for seeds produced by DH12075, Topas, Westar plants grown at CT (21/18 °C) and MT (34/18 °C). (B) Percentage of defective seedlings per categories of defects: not viable (dark blue), with one cotyledon (red), asymmetrically positioned cotyledons (green), twin seedlings (turquoise), with one cotyledon and defective root (purple), without any root (orange) and other categories (light blue). The analysis was performed on seedlings from seeds produced by DH12075, Topas, Westar plants grown at CT (21/18 °C) and MT (34/18 °C). (C-I) Range of observed seedling phenotypes. A wild-type seedling is shown in C. Seedlings (D, E) are with one cotyledon (red), Seedling in F is with two roots and two shoots (turquoise), In G, zoom in of a seedling without root (orange). In (H, I) are shown seedling from the other categories (light blue). The small coloured square in the upper right corner refers to the categories. Scale bars represent 10 mm.

In CT, the rate of defective seedlings was 2.1 % for DH12075, 5.3% for Topas and 4.3 % for Westar (Fig. 5B). In MT, the percentage of defective seedlings increased to 12 % for DH12075, 15.4 % for Topas and 16.8 % for Westar (Fig. 5B). Several types of defects were detected, from which some may partially be the consequence of the defects observed during the early embryo phenotyping analysis (Fig. 2, Figs 5C-I). We observed viable seedlings with unequal shape of cotyledons (0.67-1.1 % in CT, 1.5-2.5 % in MT) or only one cotyledon (0.44-2.7 % in CT, 3.2-10.7 % in MT, Figs 5D-E). This defect was in some cases combined with either the absence of a root or shoot apical meristem or an absence of both (Figs 5G-I). Those seedlings completely stopped growing or just delayed their development until a new root or shoot was developed, thanks to the high plasticity of plant regeneration. Other detected phenotypes were, for example, seedlings with three cotyledons and twin seedlings occurring in less than 1 % of germinated seeds (Fig. 5F).

Changes in organ growth and number indicate that high temperatures altered organogenesis. Problems with root and shoot growth point to the misregulation of the establishment of either a root or shoot apical meristem and other embryonic organs, most probably controlled by the auxin distribution during early embryo development.

#### 3.3.4 Seed metabolites content

As *Brassica napus* is an oilseed crop, the quality and composition of oil and the content of glucosinolates and nitrogen compounds were tested in seeds. Westar and DH12075 seeds from HT were only tested in one biological replicate due to the low amount of viable seeds produced. The oil content decreased by 3% in all the cultivars in MT and by 10-15 % in HT compared to CT (Fig. 6, Supplementary Table 4). The highest decrease was found in Westar cultivar from 43.38 % (CT) to 38.81 % (MT) and 26.97 % (HT) in dry matter (Fig. 6C, Supplementary Table 4). Changes were also detected in oil quality. The main compound in rapeseed oil is oleic acid, which was increased by 3 % in MT at the expense of desaturated linoleic acid and linolenic acid, which decreased by 0.5 % and 1 %, respectively. In HT, on the contrary, the content of oleic acid dropped, and the levels of linoleic acid and linolenic acid increased. Contents of palmitic, stearic, and erucic acid in rapeseed oil are low (< 5 %), and their changes are not biologically relevant (Fig. 6).

**Fig. 6.**
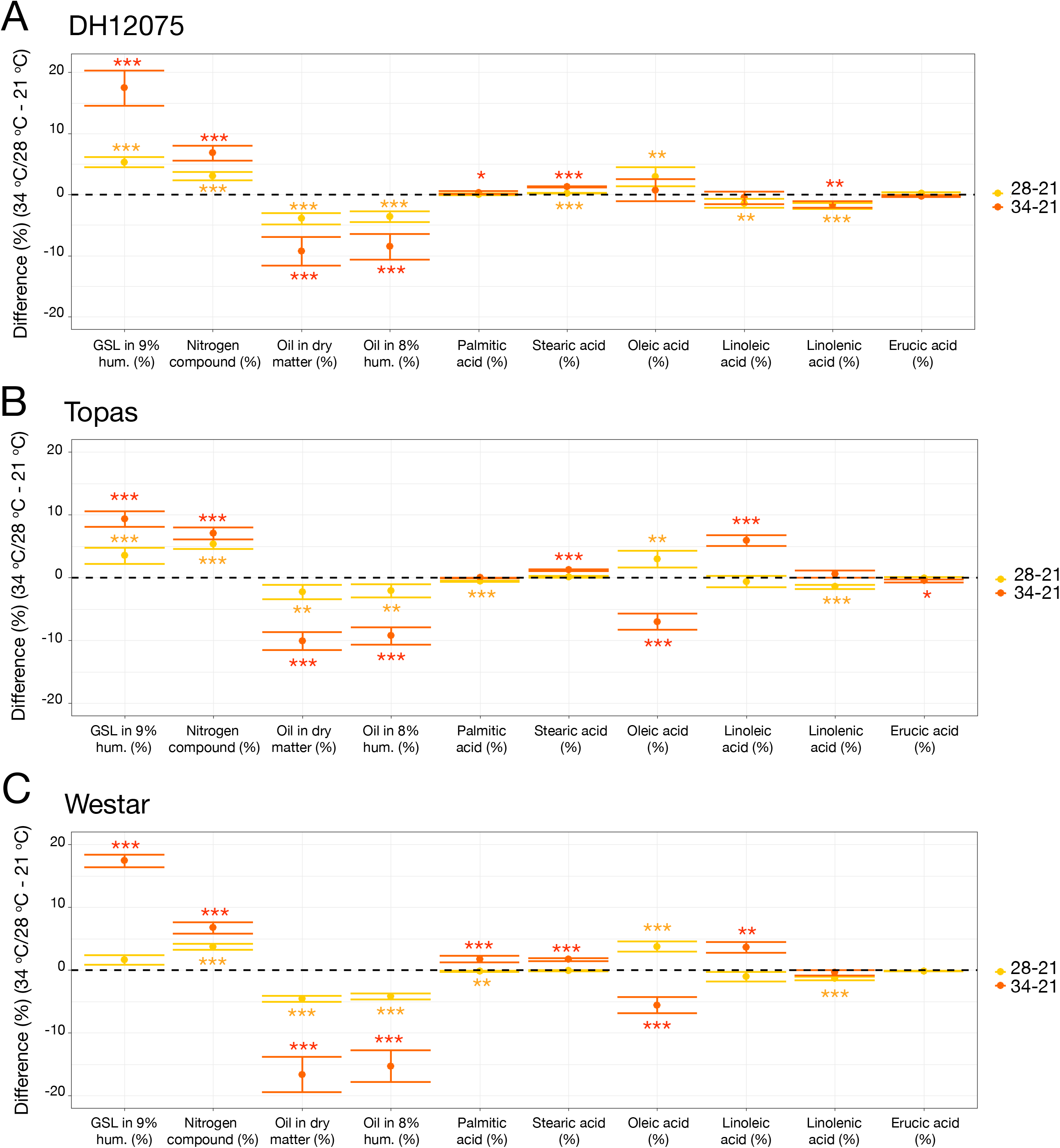
Differences in Glucosinolates, Nitrogen and seed oil levels induced by high temperatures. Graphs are displaying the differences between MT and CT and between HT and CT in levels of glucosinolates (GSL in 9% humidity, μmol/g), Nitrogen compound (%), total oil in dry matter (%) and in 8% humidity (%), palmitic acid (% of total oil), stearic acid (% of total oil), oleic acid (% of total oil), linoleic acid (% of total oil), linolenic acid (% of total oil) and erucic acid (% of total oil). The dots represent the mean difference between the temperatures and error bars, the 95 % confidence interval. Asterisks indicate statistically significant difference in MT (yellow) and HT (orange) in a paired Student’s t-test (t-test; *, **, and *** correspond to P-values of 0.05 > p > 0.01, 0.01 > p > 0.001, and p < 0.001, respectively). The measurements were done in seeds of DH12075 (A), Topas (B), and Westar (C). Source data are found in Supplementary Table S4. The measurements were done in two biological replicates, three technical replicates, except for HT seeds of DH12075 and Westar due to the reduced number of seeds produced at HT.

Seeds also contain higher levels of nitrogen compounds by more than 3 % in MT and more than 6 % in HT (Fig. 6, Supplementary Table 4). Content of glucosinolates was increased by 1.5-6 μmol/g in MT and 9-18 μmol/g in HT. In total numbers, Topas exhibits the lowest increase of glucosinolates (19.75 μmol/g in HT). On the other hand, in DH12075, the level of glucosinolates increased to 29.82 μmol/g, which can impair the nutritional properties of the seeds (Fig. 6).

### 3.4 Hormonal and transcriptional changes

The changes described above suggest that warm temperatures alter plant growth, the establishment of new organs, and the timing of their development. These processes are controlled by the biosynthesis, transport, signalling and degradation of phytohormones. To better understand how warm temperatures would unbalance the phytohormone homeostasis, auxin and ABA precursors and metabolites were quantified, and the expression of key enzymes was investigated.

#### 3.4.1 Auxin metabolism

Auxin metabolites content was tested in pistils, 5 DAP seeds and 26 DAP seeds from CT and HT in Westar. Increased levels of tryptophan (TRP), indole-3-acetamide (IAM) and indole-3-acetonitrile (IAN) were found in pistils without any changes in indole-3-acetic acid (IAA) and degradation metabolites levels except for decreased levels of 2-oxoindole-3-acetyl-1-O-ß-d-glucose (oxIAA-Glc) (Fig. 7, Supplementary Table S5). Expression of *MYB34*, encoding an MYB transcription factor regulating the expression of *CYP79B2/3* enzymes (Celenza *et al*., 2005), was significantly increased in HT pistils and leaves (Fig. 8A). The upregulation of *CYP79B2/3* in HT pistils and leaves is not considered significant (Fig. 8A). Regarding auxin degradation, only *GRETCHEN HAGEN3.5* (*GH3.5*) is significantly upregulated, even though the amino-acid conjugate levels remain unchanged. Also, the *DAO* genes expression was down-regulated, correlating with the decreased levels of oxIAA-Glc (Fig. 7, Fig. 8B). We did not see any phenotypic changes of non-pollinated pistils and the auxin homeostasis seems to be maintained in HT conditions. Therefore, the increased levels of TRP, IAM and IAN may be related to secondary metabolites protection rather than auxin biosynthesis.

**Fig. 7.**
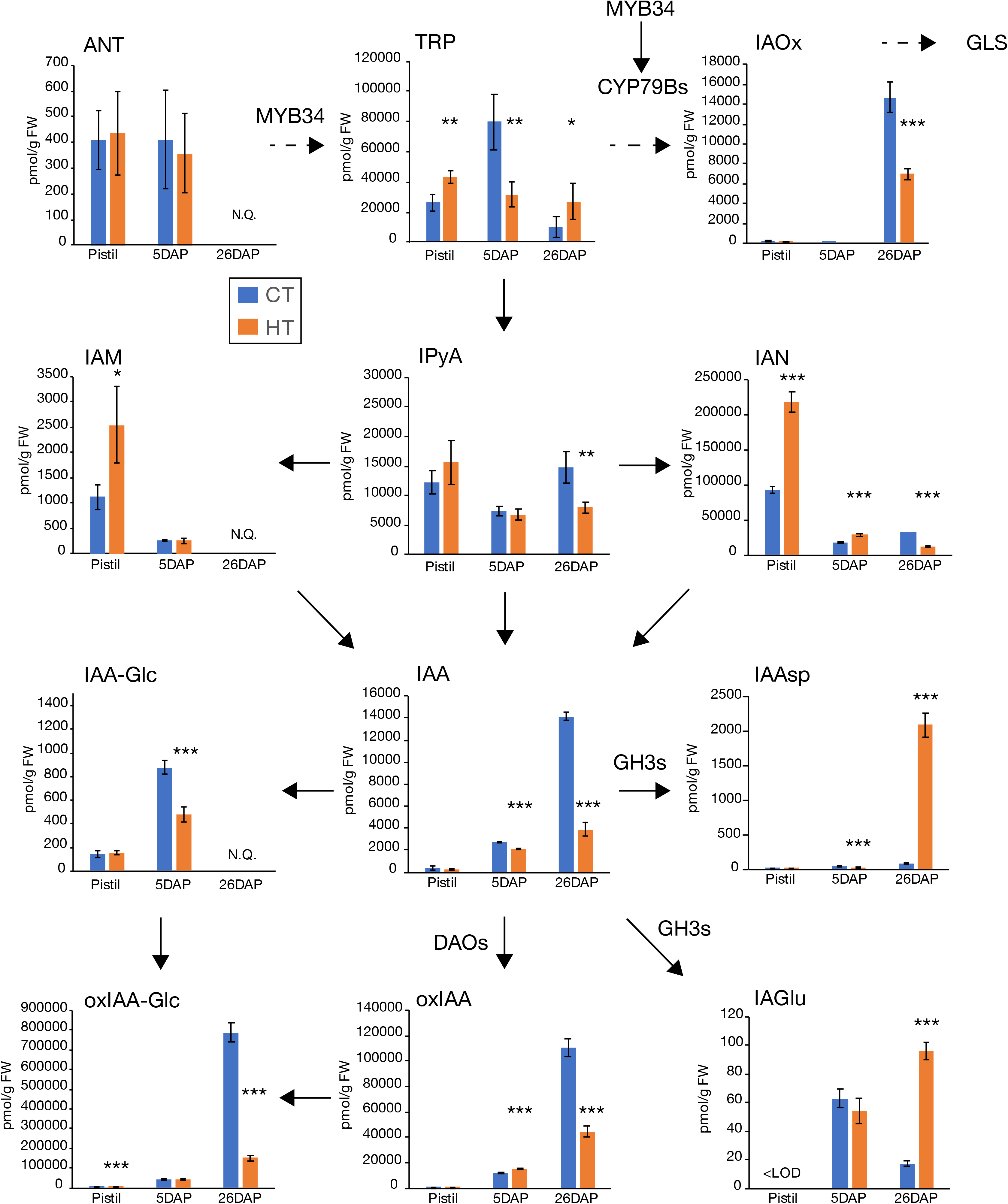
Differences in auxin and auxin metabolites levels induced by high temperatures in Westar pistils, 5 DAP and 26 DAP seeds. Graphs displaying the levels (pmol/g FW) of anthranilate (ANT), tryptophan (TRP), indole-3-acetaldoxime (IAOx), indole-3-pyruvic acid (IPyA), indole-3-acetamide (IAM), indole-3-acetonitrile (IAN), indole-3-acetic acid (IAA), IAA-glucose (IAA-Glc), IAA-aspartate (IAAsp), IAA-glutamate (IAGlu), 2-oxoindole-3-acetic acid (oxIAA) and oxIAA-glucose (oxIAA-Glc). Measurements were done in two biological replicates in five technical replicates in Westar pistils, 5 DAP seeds and 26 DAP seeds from plants grown at CT (blue) and HT (orange). Shown is the average ± S.D. of one of the two biological replicates. NQ: not quantified; <LOD: below the limit of detection. Asterisks indicate statistically significant difference in HT in a paired Student’s t-test (t-test; *, **, and *** correspond to P-values of 0.05 > p > 0.01, 0.01 > p > 0.001, and p < 0.001, respectively). Arrows indicate the direction of the biosynthesis pathway (plain arrow, direct reaction; dashed arrow, multiple-step enzymatic reactions). The names closed to the arrows indicate the enzymes involved in the reactions and whose gene expression was tested in Fig. 8. Source data are shown in Supplementary Table S5.

**Fig. 8.**
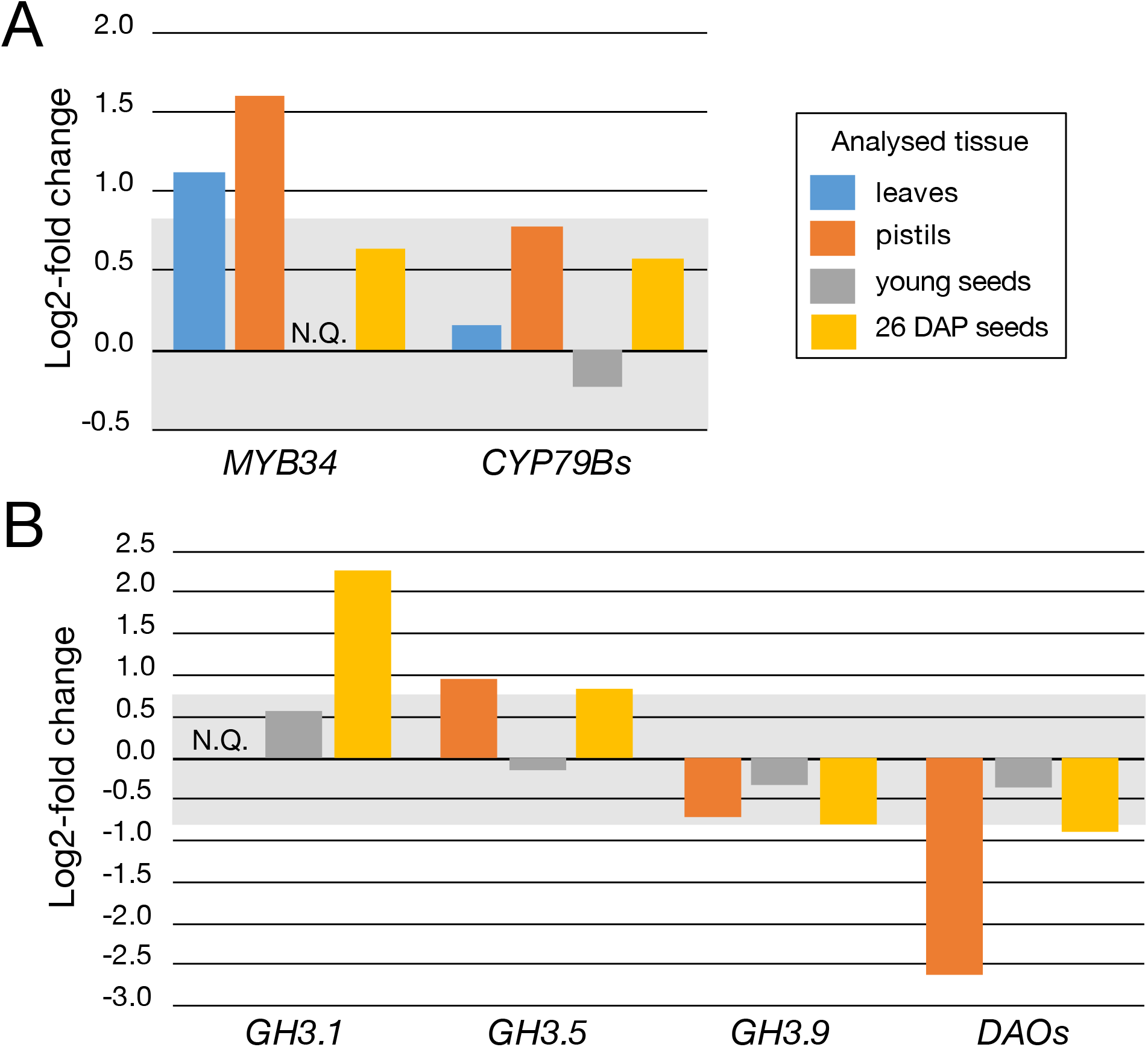
Expression analysis of genes involved in auxin homeostasis in Westar leaves, pistils, young seeds and 26 DAP seeds. (A) Expression analysis by RT-qPCR of *BnaMYB4* and *BnaCYP79Bs* in Westar leaves (blue), pistils (orange), young seeds (grey) and 26 DAP seeds (yellow). (B) Expression analysis by RT-qPCR of *BnaGH3.1*, *BnaGH3.5, BnaGH3.9* and *BnaDAOs* in Westar leaves, pistils, young seeds and 26 DAP seeds. The graphs are displaying the log2-fold changes in expression between CT and HT. The grey zone covers the [−0.8 to 0.8] log2-fold changes, considered as non-significant changes. Primers and LOC information are presented n Supplemental Table S1. N.Q.: not quantified.

In 5 DAP seeds, a decrease of TRP, IAA, and some degradation metabolites was detected, indicating altered auxin homeostasis in HT (Fig. 7). However, the expression of *GH3s*, *CYP79Bs,* and *DAOs* was not significantly changed in seeds bearing EG embryos in HT (Fig. 8B). IAA is essential for regulating embryonic morphogenesis with the establishment of an apical-basal polarity axis (Benková *et al.*, 2003; Friml *et al.*, 2003), endosperm development (Figueiredo *et al.*, 2015), and seed coat development (Figueiredo *et al.*, 2016). Decreased IAA levels in HT may alter the synchronous development of the embryo, endosperm and seed coat, therefore explaining the higher abortion rate. Also, the observed embryonic defects (e.g., altered morphology, Fig. 2) may be explained by lower auxin levels (Robert *et al.*, 2013). In CT, 5 DAP old seeds contain embryos at 2-cell or 8-cell stages (Table 2), which requires the transport of auxin from the seed integuments for its proper development (Robert *et al.*, 2018). In HT, 5 DAP old seeds bear globular embryos, which produce their auxin needed for its morphogenetic development (Robert *et al.*, 2013).

In 26 DAP seeds, higher TRP content with lower levels of direct auxin precursors (indole-3-pyruvic acid (IPyA), indole-3-acetaldoxime (IAOx) and IAN) was observed in HT seeds (Fig. 7). IAA levels are also decreased in 26 DAP seeds. Furthermore, a shift from the oxidative auxin degradation by the DAO enzymes toward the production of amino acid-auxin conjugates was observed (less 2-oxoindole-3-acetic acid, oxIAA, and oxIAA-Glc, more indole-3-acetic acid-aspartate, IAAsp, and indole-3-acetic acid-glutamate, IAGlu). This was confirmed by an expression analysis showing downregulation of *DAO*s genes and upregulation of some *GH3* genes (Fig. 8B). Surprisingly, the decrease of IAN levels was detected in this tissue even though elevated levels of glucosinolates (Figs 6 and 7) were found in mature seeds.

#### 3.4.2 ABA biosynthesis

The total concentration of ABA was analysed in the same tissues as for auxin. Recently, it has been shown that an accumulation of IAM in plant tissue impairs growth and seed development and represses temperature stress-related genes (*HSFA2*, *HSFA3*) (Sánchez-Parra *et al.*, 2021). Moreover, the crosstalk between the levels of IAM (and AMIDASE1 activity) and ABA has been identified (Pérez-Alonso *et al.*, 2021). In HT pistils, we found higher levels of IAM, reduced expression of ABA biosynthetic genes *NCED9* and significant lower ABA content (Figs 4C,D, Fig. 7, Supplementary Table S6). It suggests reduced protection of pistils to temperature stress, eventually leading to a reduced fertilization rate.

No changes in ABA levels were detected in 5 DAP seeds. As one would expect an increase of ABA linked to the applied temperature stress, the steady ABA levels may relate to the abortion of seeds due to lower stress protection (Fig. 4C, Supplementary Table S6).

Twenty-six DAP seeds are in the seed maturation stage, preparing for dormancy. ABA is known to promote dormancy in many crops. The significant decrease of ABA levels in 26 DAP seeds from HT indicates that HT limits seed dormancy progression by reducing the *NCED9* expression, as observed in our qPCR experiments (Figs 4C,D, Supplementary Table S6). However, no expression changes were identified for ABA signalling when we tested *BnaABI1* expression in pistils and 26 DAP seeds, where ABA levels were reduced (Figs 4C,D). Lower ABA levels may explain the PHS phenotype (Figs 4A-B, Table 3) when the embryo continues to grow, breaks the seed coat, and finally desiccates during fruit ripening. These seeds are not viable. Because they did not progress into the seed maturation phase, their oil content would be decreased. Therefore, HT would impair the production of high-quality seeds and reduce the final yield for food production.

#### 3.4.2 Transcriptional changes in sugar, stress and photoperiod signalling pathways

Growth at high temperatures (HT) altered hormonal levels and metabolites content in 26 DAP seeds. We analysed whether HT would alter the expression in chosen genes involved in the spliceosome, sugar metabolism and stress response. The gene encoding fructose 1,6-biphosphate aldolase 6 (FBA6) enzyme was investigated because of its putative relation in the crosstalk between sugar signalling, photoperiod and stress signalling pathways. DORMANCY/AUXIN ASSOCIATED FAMILY PROTEIN 1 (DRM1) and DRM2 are often used as markers for dormancy release due to their expression in the dormant stages of meristematic tissues, including during seed dormancy (Rae *et al.*, 2014). We found that *BnaFBA6* and *BnaDRM2* were significantly upregulated at HT in 26 DAP seeds (Fig. 4E). Studies showed that *FBA6* is upregulated in seedlings in response to sugar treatments and heat stress in Arabidopsis (Lu *et al.*, 2012), which might correlate with the expected increased sugar metabolism and partial loss of dormancy in HT 26 DAP seeds showing PHS phenotype (for which DRM2 may be the marker). The more specific role of DRM2 in the temperature response during seed maturation and dormancy would require to be elucidated.

Recent studies indicated that misregulation of the spliceosome complex leads to defective development of the male and female gametophytes resulting in embryo lethality in Arabidopsis (Kulichová *et al*., 2020; Slane *et al*., 2020). Both high temperatures and altered ABA levels in seeds may regulate the levels of the spliceosome components (Cruz *et al.*, 2014; Jiang *et al.*, 2017). As proper splicing of the pre-mRNA is required for the function of many of the essential genes, we have checked the expression of three genes involved in the spliceosome complex *SLU7*, *SCL30A* and *HSP70* (annotated in Genebank as the subunit 37c of the RNA polymerase II, *RNPII37C*). *SLU7* and *SCL30A* are significantly upregulated in the young and 26 DAP seeds developed at HT, while *RNPII30C* is upregulated in young seeds and downregulated in 26 DAP seeds (Figs 2I). Misregulation of the spliceosome activity in seeds at HT may be connected with the decreased fertilization rate and increased defects in embryo and seed development.

## 4. Discussion

In this study, phenotypic responses to high temperatures were described on the whole flowering plant level. Plants reacted to HT during reproductive development by prolonging the growth of the main stem and their flowering time and developing more branches to compensate for the lower pollination and fertilization rates and increased seed abortion. Thus, based on the prolonged duration of flowering, we hypothesized an alteration in the photoperiod or maturation-related regulation of hormonal levels during plant development. Moreover, increased branching may be associated with decreased apical dominance. As previous studies applied heat stress for limited periods, it hardly showed the changes on the whole plant level, although partial hints in accordance with our results were published. Indeed, HT increased plant growth and their above-ground biomass regardless of the genotype (Chen *et al.*, 2020a). And significantly higher production of lateral inflorescences led to higher production after stress release and return to control conditions (Young *et al.*, 2004). Both studies showed that these changes are not specific to the cultivars used in our study but could be broadly applied.

We did not find any biologically significant changes in pollen viability, which could cause a decrease in fertilization rate and seed number per pod. These findings do not correspond with published data of reduced pollen viability after 7-day heat treatment in *B. napus* and *B. rapa* (Chen *et al.*, 2020a; Young *et al.*, 2004; Annisa *et al.*, 2013). Our growth setup includes lower night temperatures (18 °C in all conditions) and ramping up and down to stress temperatures during the day. The pollen development may benefit from the colder night temperatures to cope with the stress conditions.

A significant decrease in fertilization rate (occurrence of unfertilized ovules) may be caused by the infertility of the ovules, lower pollen germination (Young *et al.*, 2004), or lower transmission of pollen tubes through the transmitting tract. The percentage of aborted seeds increased with increasing temperatures, which was also noticed without further investigations (Chen *et al*., 2020a). In our study, only DH12075 had a very low abortion rate for all tested temperatures.

To our knowledge, the dynamics of embryo development in higher temperatures have not yet been studied. HT accelerates early embryo development, which may cause the misregulation of essential signalling pathways. The described embryonic defects indicate irregularities in cell division patterns in HT compared to CT. Defective embryos present imprecise determination of cell fate in suspensor cells and misregulation in organ formation and determination in the proembryo. This defective development affects 15-40 % of embryos in HT. Similar defects were previously described in Arabidopsis mutants in auxin biosynthesis, perception, and signalling pathway. Extra horizontal and vertical divisions of suspensor and missing cotyledons were described in *tir1afb235* quadruple mutant (Prigge *et al.*, 2020). Suspensor cell identity was disturbed in *iaa10* mutant leading to missing root tissue (Rademacher *et al.*, 2012) or forming secondary embryos (Radoeva *et al.*, 2019). Maintenance of both embryonic and suspensor cell fate is essential for correct development. We hypothesize that HT alters auxin distribution and/or signalling in both suspensor and embryo proper, which results in the described embryonic defects. Some of the defects were related to phenotypes observed in germinated seedlings, so the embryos were viable and went through the seed maturation phase. Other defects increased the seed abortion penetrance during later stages of seed development.

HT treatment caused a decrease in the number of viable seeds per silique: < 5 seeds for DH12075 and Westar, and < 10 seeds for Topas. The reduction of seed number was combined with a reduction in the seed weight. A comparable effect was observed in rice (Wu *et al.*, 2016). Angadi *et al.* (2000) suggested that the optimal daytime temperature for *B. napus* is closer to 28 °C instead of 20 °C or 35 °C (Angadi *et al*., 2000). In our study, MT induced only a longer main stem but not the development of primary branches. As we did not study the growth and production of branches, we cannot evaluate if the lost in seed number per silique, seed weight, and altered oil composition observed in the main stem could be compensated by increasing total flower production.

The reduced quality of the harvest was partially caused by premature seed germination (sprouting) initiated by HT. It was noticed in all three studied cultivars, and most severely in Topas (30.4-73.4 %). Seed sprouting and lower ABA content in HT than CT indicate that seeds did not progress into the dormancy phase but continue directly to germination. Primary dormancy in *B. napus* reaches a peak of ABA concentration between 30 to 40 DAP and decrease rapidly to 0-15 % in the next 2 to 3 weeks (Huang *et al.*, 2016). Therefore, our experiment showed that the embryos’ accelerated development caused seed sprouting before the expected ABA accumulation. The dormancy induction in developing seeds is achieved by embryonic ABA production (Groot and Karssen, 1992). The production in maternal tissue can only partially induce dormancy in Arabidopsis in the absence of zygotic ABA (Kanno *et al.*, 2010). *NCED6* and *NCED9* are the main ABA biosynthetic genes expressed preferentially in developing seeds (Lefebvre *et al*., 2006). Disruption of their activity reduced seed dormancy in *Arabidopsis* (Lefebvre *et al*., 2006). Our expression analysis identified a correlation between the reduced *NCED9* expression levels and the reduced ABA content in 26 DAP seeds at HT. Whether HT affects the site of production, transport, reception or degradation of ABA in seeds remains to be elucidated.

HT treatment caused fine-tuning changes in auxin biosynthesis and degradation while reducing auxin levels in 26 DAP seeds. Levels of oxIAA are decreased by half (even more for oxIAA-Glc), which are compensated by an increase of conjugative degradation by GH3 proteins, a phenomenon which has also been described in the *Arabidopsis dao1-1* mutant line (Mellor *et al.*, 2016). Auxin has been shown to act upstream of ABA in controlling seed dormancy (Liu *et al.*, 2013). Auxin signalling controls the expression of ABA signalling genes involved in the progression and maintenance of seed dormancy. Therefore, in 26 DAP seeds at HT, seed sprouting may be induced by reduced ABA levels and a reduced auxin input on ABA signalling.

We did not observe a high reduction in the germination of fully and partially filled seeds suggesting that imperfectly developed seeds and severely defective embryos underwent abortion during development. Still, we found up to 16.8 % of defective seedlings at MT (versus a maximum of 5.3 % at CT). Most of these seedlings were viable, and despite slower growth, they progressed into normal development. The defects observed in seedlings are connected to the embryonic defects and, most probably, the changes in hormonal levels. Therefore, the misregulated pathways involved in organogenesis would benefit from further analysis.

We detected a decrease in total oil content (3-4 % in MT and 10-15 % in HT compared to CT), but an increase of 2-3 % (at 25 °C) was previously described (Aksouh-Harradj *et al.*, 2006). On the other hand, the oil quality (estimated by the ratio of oleic, linoleic and linolenic acids) was changed similarly, although not significantly in MT. The situation is different in HT, where we found no changed (DH12075) or a decrease by 5-7 % of oleic acid for the other two cultivars. In accordance, we did not find any change in the levels of erucic acid. Oil content was shown to be strongly affected by drought (decrease by 6.5-14 %). However, its effect on the fatty acid pattern was the opposite of our observations in MT and in accordance with our observations in HT (Hatzig *et al.*, 2018). These findings implicate sufficient adaptation of *B. napus* to MT. However, long-term HT could similarly affect the plants as drought stress during flowering and seed filling periods (Hatzig *et al.*, 2018).

A general tendency to higher GLS content in seeds was observed in our study for all tested cultivars, contrary to previous studies (Rao *et al.*, 2021, Aksouh-Harradj *et al.*, 2006). On the other hand, such an increase was not significant in drought stress (Hatzig *et al.*, 2018). Moreover, GLS content varies with the circadian rhythm (Huseby *et al.*, 2013), and mutants in GLS metabolism exhibited lower levels of the heat-shock stress protein 90 (HSP90) and reduced tolerance to elevated temperatures (Ludwig-Müller *et al.*, 2000). Therefore, GLS play an essential role in heat stress protection for plants, but the specific role during reproductive growth (ovules and seed development) needs to be evaluated.

Our phenotyping analysis was complemented by hormonal profiling. In general, plants mostly react to stress by increasing their ABA production to regulate their water balance, reduce desiccation, and, in the longer term, to regulate the senescence of different plant organs. Surprisingly, we found that ABA levels were decreased in non-pollinated pistils and 26 DAP seeds, while they remained unchanged in 5 DAP seeds. It may suggest that Brassica plants have little abilities to protect themselves in long-term heat stress and, therefore, led to a high abortion rate, decreased yield, and in the case of 26 DAP seeds, reduced dormancy. These results contrast with research done on pea seeds (Kaur *et al.*, 2020) and rice panicles (Wu *et al.*, 2016).

In summary, elevated temperature influences the reproductive growth of *B. napus* in different manners. A decrease in seed yield is compensated by the development of primary branches and prolonged growth of the main stem. Seed development is negatively affected, leading to a higher abortion rate, most probably due to misregulated embryo development and misregulated auxin signalling. HT changed phytohormone levels, especially ABA, which we believe to be the reason for higher PHS. Harvested seeds showed reduced quality, in particular lower oil content and higher GSL content. Our observations pinpointed several aspects that will require more in-depth analysis to understand seed development in response to suboptimal growth temperatures.

## Abbreviations in alphabetical order

ANT: anthranilate
ABA: abscisic acid
CT: control temperature
DAP: days after pollination
DAV: days after vernalization
EG: early globular
FT: flowering time
GLS: glucosinolates
HS: heat stress
HSF: heat stress transcription factor
HSP: heat shock protein
HT: high temperature
IAA: indole-3-acetic acid
IAAsp: indole-3-acetic acid-aspartate
IAA-Glc: indole-3-acetyl-1-O-ß-d-glucose
IAGlu: indole-3-acetic acid-glutamate
IAM: indole-3-acetamide
IAN: indole-3-acetonitrile
IAOx: indole-3-acetaldoxime
IPyA: indole-3-pyruvic acid
LG: late globular
LMS: length of main flowering stem
LOD: limit of detection
MG: mid-globular
MT: mid-temperature
NF: number of flowers on the main stem
NL: number of leaves
NB: number of primary branches
NF: number of flowers on the main stem
NO: number of ovules
NQ: Not quantified
NS: number of seeds
oxIAA: 2-oxoindole-3-acetic acid
oxIAA-Glc: 2-oxoindole-3-acetyl-1-O-ß-d-glucose
PHS: preharvest sprouting
RGR: relative growth rate
RSD: relative standard deviation
SL: silique length
TRP: tryptophan

## Supplementary data

Supplementary data are available at *JXB* online.

*Table S1*. Primers used in qPCR analysis and LOC number of amplified genes.

*Table S2*. Pearson correlation coefficient between the length of the main inflorescence stem, the flowering time duration, and the number of flowers.

*Table S3.* Pearson correlation coefficient between the siliques growth rate, the seed number per silique, and the growth temperature.

*Table S4*. Glucosinolates, Nitrogen, and seed oil measurement (source data of Fig. 6).

*Table S5.* Auxin and auxin metabolites measurements (source data of Fig. 7).

*Table S6.* ABA measurements (source data of Figure 4).

*Fig. S1.* Setup of growth temperatures in the greenhouse chambers.

*Fig. S2.* High temperatures affect the growth parameters of Brassica flowering plants.

*Fig. S3.* The number of ovules is unchanged, but embryonic development is accelerated when plants are grown at high temperatures.

*Fig. S4*. Pollen grain development is not affected by our stress growth conditions.

*Fig. S5*. High temperatures affect embryo development. Original pictures presented in Fig. 2.

*Fig. S6.* Silique growth rate is affected at higher temperatures.

## Acknowledgements

The authors thank Jarmila Greplová, Kamila Wisnerová and Michaela Mrvková for their help with phytohormone analyses. We acknowledge the Core Facility CELLIM supported by the Czech-BioImaging large RI project (LM2018129 funded by MEYS CR) and the Core Facility Plant Sciences of CEITEC Masaryk University for their support with obtaining scientific data presented in this paper. This work was supported by the Czech Science Foundation (project no. 19-05200S) to HSR, from the Ministry of Education, Youth and Sports of the Czech Republic with the European Regional Development Fund-Project “SINGING PLANT” (No. CZ.02.1.01/0.0/0.0/16_026/0008446) to HSR.

## Author contributions

KM: investigation (all), formal analysis, visualization, writing: original draft preparation, writing: review and editing

UP: investigation (phenotyping, RT-qPCR), writing: review and editing

MS: investigation (phenotyping, RT-qPCR), writing: review and editing

IS: formal analysis (statistical analysis), visualization

AP: investigation (hormone profiling)

LE: investigation (GLS, nitrogen and oil measurements)

ON: investigation (hormone profiling), funding acquisition

HSR: conceptualization, funding acquisition, project administration, supervision, visualization, writing: original draft preparation, writing: review and editing

## Data availability statement

The data supporting the findings of this study are available within the paper and within its supplementary materials published online. A request may be made to the corresponding author, H.S. Robert.

