## Supplementary material for "Effects of long-term high-temperature stress on reproductive growth and seed development in development in *Brassica napus*"

Supplementary data are available at *JXB* online.
*Table S1*. Primers used in qPCR analysis and LOC number of amplified genes.
*Table S2*. Pearson correlation coefficient between the length of the main inflorescence stem, the flowering time duration and the number of flowers.
*Table S3.* Pearson correlation coefficient between the siliques growth rate, the seed number per silique and the growth temperatures.
*Table S4*. Glucosinolates, Nitrogen and seed oil measurement (source data of Fig. 6).
*Table S5.* Auxin and auxin metabolites measurements (source data of Fig. 7).
*Table S6.* ABA measurements (source data of Figure 4) .
*Fig. S1.* Setup of growth temperatures in greenhouse chambers.
*Fig. S2.* High temperatures affect growth parameters of Brassica flowering plants.
*Fig. S3.* The number of ovules is unchanged, but the development of the embryo is accelerated when plants are grown at high temperatures.
*Fig. S4*. Pollen grain development is not affected by our stress growth conditions.
*Fig. S5*. High temperatures affect embryo development. Original pictures presented in Fig. 2.
*Fig. S6.* Silique growth rate is reduced at higher temperatures.

*Supplementary Table S1*. Primers used in qPCR analysis and LOC number of amplified genes

| Primer name | Primer sequence | Targeted LOCs in *B. napus* | Homologous genes in *Arabidopsis* |
|---|---|---|---|
| BnaMYB34-FW<br>BnaMYB34 REV | ACGTCGATTCTCCGACCAA<br>TATGAAACCGCTTGACGCTG | LOC106374996<br>LOC106434612 | *MYB34*<br>*At5g60890* |
| BnaCYP79B-FW<br>BnaCYP79B-REV | CTTGGAAGTTGCCTGAGAATG<br>GAACGATGCTCTGGGAGTC | LOC106439341 | *CYP79B2*<br>*At4g39950*<br>*CYP79B3*<br>*At2g22330* |
| BnaDAOs-FW<br>BnaDAOs-REV | TTTGTGGATGCTGAACATCCG<br>TAAGCTTGAGAGCTTCTCCATC | LOC106414697<br>LOC106430970<br>LOC106369131<br>LOC106399859<br>LOC106380301<br>LOC106382334<br>LOC106414671<br>LOC106369133<br>LOC106362101<br>LOC106379949 | *DAO1*<br>*At1g14130*<br>*DAO2*<br>*At1g14120* |
| BnaGH3.9s-FW<br>BnaGH3.9s-REV | ACGTGGTTCTAAGCATCGAC<br>GCGAGGAATGCCTTGTTTTC | LOC106392343<br>LOC106396677<br>LOC106392367<br>LOC106450413 | *GH3.9*<br>*At2g47750* |
| BnaACGH3.1-FW<br>BnaACGH3.1-REV | AGTTGTTAGTGAGGGACGGA<br>CGGTAAACCGAGTTCAACGA | LOC106409452<br>LOC106432954 | *GH3.1*<br>*At2g14960* |
| BnaGH3.5s_FW<br>BnaGH3.5s_REV | AAGAACGCAATGACACACCT<br>TGGCCAGGGATAGAACTTGT | LOC106409348<br>LOC106376087<br>LOC106440950<br>LOC106445777 | *GH3.5*<br>*At4g27260* |
| BnaNCED9-FW2<br>BnaNCED9-REV2 | GTGACGGTAAGTTCGGAGGA<br>GATCTCACATTTTCCTCGTCGT | LOC106354557<br>LOC106354700<br>LOC106441091 | *NCED9*<br>*At1g78390* |
| BnaABI1-2 FW1<br>BnaABI1-2 REV1 | TTCGGACAAGAAGGCGG<br>AATTCACCGTCGGTTTACTCAA | LOC106390346 | *ABI1*<br>*At4g26080* |
| BnaPHYA-FW1<br>BnaPHYA-REV1 | TTGTGTCAGAAGCAGCTCAG<br>TCCAGATCCAAGCACCCTTC | LOC106400740<br>LOC106346267<br>LOC106417700<br>LOC106420085 | *PHYA*<br>*At1g09570* |
| BnaELF4-FW1<br>BnaELF4-REV1 | GGAGCAGGGAGGAGAAGATC<br>GGTGATTGTCGTTGACCTGC | LOC106420605<br>LOC106394477<br>LOC106394476<br>LOC106438048<br>LOC106399222<br>LOC106454592<br>LOC106399325 | *ELF4*<br>*At2g40080* |
| BnaRNPII37c-FW1<br>BnaRNPII37c-REV1 | TAACGACAAGGGAAGGCTGT<br>GTGTTCCTCATCCTCTGCCT | LOC106377472<br>LOC106353095 | *HSP70*<br>*At3g12580* |
| BnaSCL30A-FW2<br>BnaSCL30A-REV2 | AACGATCAAGGGGAAGGACA<br>CCTGTAGACTGGCGAACGT | LOC106412002<br>LOC106452514 | *SCL30A*<br>*At3g13570* |
| BnaSLU7-FW1<br>BnaSLU7-REV1 | GTCGTGGTGGAAGGATCAAC<br>CTTCCTCAGCCGCCTCTATC | LOC106446239<br>LOC106453449 | *SMP1*<br>*At1g65660* |

| | | | |
|---|---|---|---|
| BnaFBA6-FW2<br><br>BnaFBA6-REV2 | GTTGAAGACTTGGGGAGGGA<br><br>TCAGAGTTAGCCTTGCACCT | LOC106391482<br>LOC106395718<br>LOC106367801<br>LOC111206734<br>LOC106367922<br>LOC106452433<br>LOC106428297<br>LOC106445170 | *FBA6*<br>*At2g36460* |
| BnaDRM2-FW2<br><br>BnaDRM2-REV2 | TCTCACCCAAACTCTCCCAC<br><br>TCACCAGCATGTCACTCCAT | LOC106451187<br>LOC106347199<br>LOC106437808 | *DRM2*<br>*At2g33830* |
| BnaTMA7-FW1<br><br>BnaTMA7-REV1 | TTCCTGTGTTTTATCCATGTAGCC<br><br>CAGTCACTCTCCTACGAACATGATAG | LOC106452133 | *TMA7*<br>*At1g15270*<br>*At3g16040* |
| BnaACT7-FW1<br><br>BnaACT7-REV1 | GAGCAGCATGAAGATCAAGGT<br><br>CTTCGAGATCCACATCTGTTGG | LOC106382989<br>LOC106426760<br>LOC106426759 | *ACT7*<br>*At5g09810* |
| BnaeIF5A-FW1<br><br>BnaeIF5A-REV1 | ATCTCAGCGCTCTGATGAAGA<br><br>ACTATTGGTTTACTTGCCACC | LOC106389545 | *eIF5A*<br>*At1g13950*<br>*At1g26630*<br>*At1g69410* |

*Supplementary Table S2*. Pearson correlation coefficient between the length of the main inflorescence stem, the flowering time duration and the number of flowers.

| | LMSxFT | | | LMSxNF | | | FTxNF | | |
|---|---|---|---|---|---|---|---|---|---|
| | 21 | 28 | 34 | 21 | 28 | 34 | 21 | 28 | 34 |
| DH12075 | 0.88 | 0.63 | 0.78 | 0.77 | 0.43 | 0.77 | 0.76 | 0.57 | 0.23 |
| Topas | 0.80 | 0.79 | 0.41 | 0.81 | 0.90 | 0.44 | 0.79 | 0.77 | 0.37 |
| Westar | 0.81 | 0.63 | 0.60 | 0.81 | 0.74 | 0.22 | 0.76 | 0.58 | 0.22 |

Strong correlation 1-0.5, medium 0.5 -0.3, small < 0.3

*Supplementary Table S3.* Pearson correlation coefficient between the siliques growth rate, the seed number per silique and the growth temperatures

| Cultivars | CT (21ºC) | MT (28ºC) | HT (34ºC) |
|---|---|---|---|
| DH12075 | 0.83 | 0.96 | 0.99 |
| Topas | 0.63 | 0.92 | 0.92 |
| Westar | 0.67 | 0.92 | 0.96 |

Strong correlation 1-0.5, medium 0.5 -0.3, small < 0.3

*Supplementary Table S4*. Glucosinolates, Nitrogen and seed oil measurement (source data of Fig. 6).

| sample | | av. | | S.D. | av. | | S.D. | av. | | S.D. |
|---|---|---|---|---|---|---|---|---|---|---|
| **Cult.** | **temp** | **GSL in 9% humidity** | | | **Nitrogen compound** | | | **Oil in dry matter** | | |
| **Topas** | **21** | 10.43 | ± | 0.30 | 21.42 | ± | 0.57 | 41.36 | ± | 0.80 |
| | **CT** | RSD | | 2.86% | RSD | | 2.67% | RSD | | 1.93% |
| | **28** | 13.92 | ± | 1.52 | 26.78 | ± | 0.84 | 39.12 | ± | 0.93 |
| | **MT** | RSD | | 10.88% | RSD | | 3.12% | RSD | | 2.38% |
| | | *** | / | 0.00049 | *** | / | 0.00000 | ** | / | 0.00119 |
| | **34** | 19.75 | ± | 1.37 | 28.50 | ± | 1.45 | 31.28 | ± | 1.90 |
| | **HT** | RSD | | 6.95% | RSD | | 5.10% | RSD | | 6.07% |
| | | *** | / | 0.00000 | *** | / | 0.00000 | *** | / | 0.00000 |
| **Westar** | **21** | 8.38 | ± | 1.20 | 20.89 | ± | 0.62 | 43.38 | ± | 1.18 |
| | **CT** | RSD | | 14.33% | RSD | | 2.96% | RSD | | 2.72% |
| | **28** | 10.03 | ± | 1.17 | 24.61 | ± | 0.58 | 38.81 | ± | 0.55 |
| | **MT** | RSD | | 11.72% | RSD | | 2.36% | RSD | | 1.41% |
| | | - | / | 0.05311 | *** | / | 0.00000 | *** | / | 0.00001 |
| | **34** | 25.82 | ± | 0.42 | 27.52 | ± | 1.24 | 26.97 | ± | 3.73 |
| | **HT** | RSD | | 1.63% | RSD | | 4.50% | RSD | | 13.84% |
| | | *** | / | 0.00000 | *** | / | 0.00001 | *** | / | 0.00002 |
| **DH12075** | **21** | 11.76 | ± | 0.68 | 20.48 | ± | 0.70 | 41.02 | ± | 0.95 |
| | **CT** | RSD | | 5.77% | RSD | | 3.44% | RSD | | 2.32% |
| | **28** | 17.10 | ± | 1.44 | 23.52 | ± | 0.64 | 37.10 | ± | 1.07 |
| | **MT** | RSD | | 8.42% | RSD | | 2.73% | RSD | | 2.87% |
| | | *** | / | 0.00475 | *** | / | 0.00001 | *** | / | 0.00005 |
| | **34** | 29.82 | ± | 3.64 | 27.79 | ± | 1.60 | 31.07 | ± | 3.50 |
| | **HT** | RSD | | 12.21% | RSD | | 5.76% | RSD | | 11.26% |
| | | *** | / | 0.00002 | *** | / | 0.00002 | *** | / | 0.00023 |

| sample | | av. | | S.D. | av. | | S.D. | av. | | S.D. |
|---|---|---|---|---|---|---|---|---|---|---|
| **Cult.** | **temp** | **Oil in 8% hum.** | | | **Palmitic acid** | | | **Stearic acid** | | |
| **Topas** | **21** | 38.05 | ± | 0.74 | 4.35 | ± | 0.0§3 | 2.52 | ± | 0.10 |
| | **CT** | RSD | | 1.94% | RSD | | 0.72% | RSD | | 3.88% |
| | **28** | 35.99 | ± | 0.86 | 3.82 | ± | 0.12 | 2.68 | ± | 0.15 |
| | **MT** | RSD | | 2.38% | RSD | | 3.03% | RSD | | 5.60% |
| | | ** | / | 0.00120 | *** | / | 0.00000 | - | / | 0.06474 |
| | **34** | 28.78 | ± | 1.75 | 4.32 | ± | 0.08 | 3.76 | ± | 0.20 |
| | **HT** | RSD | | 6.07% | RSD | | 1.80% | RSD | | 5.23% |
| | | *** | / | 0.00000 | - | / | 0.51043 | *** | / | 0.00000 |
| **Westar** | **21** | 39.91 | ± | 1.09 | 4.13 | ± | 0.08 | 2.38 | ± | 0.05 |
| | **CT** | RSD | | 2.72% | RSD | | 1.84% | RSD | | 2.04% |
| | **28** | 35.71 | ± | 0.50 | 3.96 | ± | 0.09 | 2.28 | ± | 0.16 |
| | **MT** | RSD | | 1.41% | RSD | | 2.19% | RSD | | 7.18% |
| | | *** | / | 0.00001 | ** | / | 0.00451 | - | / | 0.16873 |
| | **34** | 24.81 | ± | 3.43 | 5.92 | ± | 0.76 | 4.06 | ± | 0.35 |
| | **HT** | RSD | | 13.84% | RSD | | 12.80% | RSD | | 8.61% |
| | | *** | / | 0.00002 | *** | / | 0.00047 | *** | / | 0.00001 |
| **DH12075** | **21** | 37.74 | ± | 0.87 | 4.20 | ± | 0.09 | 2.36 | ± | 0.09 |
| | **CT** | RSD | | 2.32% | RSD | | 2.25% | RSD | | 3.63% |
| | **28** | 34.13 | ± | 0.98 | 4.27 | ± | 0.08 | 2.62 | ± | 0.05 |
| | **MT** | RSD | | 2.87% | RSD | | 1.86% | RSD | | 1.77% |
| | | *** | / | 0.00005 | - | / | 0.21433 | *** | / | 0.00005 |
| | **34** | 28.58 | ± | 3.21 | 4.62 | ± | 0.31 | 3.62 | ± | 0.18 |
| | **HT** | RSD | | 11.25% | RSD | | 6.78% | RSD | | 5.06% |
| | | *** | / | 0.00023 | * | / | 0.01426 | *** | / | 0.00000 |

| sample | | av. | | S.D. | av. | | S.D. | av. | | S.D. |
|---|---|---|---|---|---|---|---|---|---|---|
| Cult. | temp | Oleic acid | | | Linoleic acid | | | Linolenic acid | | |
| Topas | 21 CT | 66.51 | ± | 0.60 | 16.96 | ± | 0.51 | 7.92 | ± | 0.38 |
| | | RSD | | 0.90% | RSD | | 3.03% | RSD | | 4.82% |
| | 28 MT | 69.50 | ± | 1.65 | 16.34 | ± | 1.00 | 6.46 | ± | 0.39 |
| | | RSD | | 2.38% | RSD | | 6.14% | RSD | | 6.09% |
| | | ** | / | 0.100196 | - | / | 0.20723 | * | / | 0.00007 |
| | 34 HT | 59.50 | ± | 1.32 | 22.85 | ± | 0.98 | 8.49 | ± | 0.66 |
| | | RSD | | 2.21% | RSD | | 4.27% | RSD | | 7.73% |
| | | *** | / | 0.00000 | *** | / | 0.00000 | - | / | 0.09582 |
| Westar | 21 CT | 70.23 | ± | 0.99 | 14.86 | ± | 0.84 | 6.28 | ± | 0.38 |
| | | RSD | | 1.40% | RSD | | 5.68% | RSD | | 6.05% |
| | 28 MT | 73.98 | ± | 0.59 | 13.85 | ± | 0.70 | 4.95 | ± | 0.25 |
| | | RSD | | 0.79% | RSD | | 5.07% | RSD | | 5.11% |
| | | *** | / | 0.00001 | * | / | 0.04761 | *** | / | 0.00003 |
| | 34 HT | 64.93 | ± | 1.47 | 18.18 | ± | 1.13 | 5.92 | ± | 0.52 |
| | | RSD | | 2.27% | RSD | | 6.20% | RSD | | 8.74% |
| | | *** | / | 0.00032 | ** | / | 0.00151 | - | / | 0.26855 |
| DH12075 | 21 CT | 68.51 | ± | 1.47 | 15.43 | ± | 0.91 | 7.03 | ± | 0.42 |
| | | RSD | | 2.14% | RSD | | 5.91% | RSD | | 5.95% |
| | 28 MT | 71.49 | ± | 0.97 | 14.02 | ± | 0.38 | 5.20. | ± | 0.26 |
| | | RSD | | 1.36% | RSD | | 2.72% | RSD | | 4.93% |
| | | ** | / | 0.00198 | ** | / | 0.00576 | *** | / | 0.00000 |
| | 34 HT | 69.08 | ± | 2.04 | 15.21 | ± | 1.06 | 5.33 | ± | 0.69 |
| | | RSD | | 2.95% | RSD | | 6.98% | RSD | | 12.85% |
| | | - | / | 0.63821 | - | / | 0.75284 | ** | / | 0.00218 |

| sample | | av. | | S.D. |
|---|---|---|---|---|
| Cult. | temp | Erucic acid | | |
| Topas | 21 CT | 0.62 | ± | 0.21 |
| | | RSD | | 33.99% |
| | 28 MT | 0.49 | ± | 0.09 |
| | | RSD | | 17.46% |
| | | - | / | 0.27948 |
| | 34 HT | 0.23 | ± | 0.08 |
| | | RSD | | 36.05% |
| | | * | / | 0.04980 |
| Westar | 21 CT | 0.52 | ± | 0.20 |
| | | RSD | | 38.64% |
| | 28 MT | 0.31 | ± | 0.14 |
| | | RSD | | 45.75% |
| | | - | / | 0.21895 |
| | 34 HT | <LOD | | |
| DH12075 | 21 CT | 1.43 | ± | 0.23 |
| | | RSD | | 15.90% |
| | 28 MT | 1.64 | ± | 0.19 |
| | | RSD | | 11.46% |
| | | - | / | 0.11958 |
| | 34 HT | 1.13 | ± | 0.04 |
| | | RSD | | 3.92% |
| | | - | / | 0,06628 |

CT: control temperature; MT: mid temperature; HT: hight temperature; av.: average; S.D.; standard deviation; RSD: relative standard deviation (ratio of the standard deviation over the average); <LOD: below the limit of detection. Asterisks indicate statistically significant difference in MT and HT compared to CT in a paired Student's t-test (t-test; *, **, and *** correspond to P-values of 0.05 > p > 0.01, 0.01 > p > 0.001, and p < 0.001, respectively). GSL is quantified as µmol/g FW. Nitrogen compound, total oil in dry matter and 8% humidity are quantified as % of FW. Fatty acids are quantified as % of total oil content.

*Supplementary Table S5*. Auxin and auxin metabolites measurements (source data of Fig. 7). One representative biological replicate is presented.

| **sample type** | **temp** | average | | S.D. | av. | | S.D. | av. | | S.D. |
|---|---|---|---|---|---|---|---|---|---|---|
| | | **TRP** | | | **IAOx** | | | **ANT** | | |
| **Pistil** | **21 CT** | 26 006.7 | ± | 5 356.7 | 194.9 | ± | 42.5 | 408.5 | ± | 113.7 |
| | | RSD | | 21% | RSD | | 22% | RSD | | 28% |
| | **34 HT** | 43 330.1 | ± | 4 248.1 | 155.8 | ± | 23.8 | 436.4 | ± | 160.6 |
| | | RSD | | 10% | RSD | | 15% | RSD | | 37% |
| | | ** | / | 0.00203 | - | / | 0.18946 | - | / | 0.78438 |
| **5 DAP** | **21 CT** | 79 755.2 | ± | 18 901.1 | 34.4 | ± | 2.5 | 410.9 | ± | 193.1 |
| | | RSD | | 24% | RSD | | 7% | RSD | | 47% |
| | **34 HT** | 31 326.1 | ± | 8 433.4 | <LOD | | | 357.4 | ± | 154.2 |
| | | RSD | | 27% | | | | RSD | | 43% |
| | | ** | / | 0.00158 | | | | - | / | 0.69579 |
| **26 DAP** | **21 CT** | 9 491.1 | ± | 7 234.0 | 14 728.9 | ± | 1 560.1 | *NQ* | | |
| | | RSD | | 76% | RSD | | 11% | | | |
| | **34 HT** | 26 807.7 | ± | 11 659.5 | 6 946.5 | ± | 541.2 | *NQ* | | |
| | | RSD | | 43% | RSD | | 8% | | | |
| | | * | / | 0.03558 | *** | / | 0.00007 | | | |

| **sample type** | **temp** | average | | S.D. | av. | | S.D. | av. | | S.D. |
|---|---|---|---|---|---|---|---|---|---|---|
| | | **IPyA** | | | **IAM** | | | **IAN** | | |
| **Pistil** | **21 CT** | 12 271.6 | ± | 2 001.4 | 1 123.2 | ± | 241.9 | 92 234.3 | ± | 4 992.8 |
| | | RSD | | 16% | RSD | | 22% | RSD | | 5% |
| | **34 HT** | 15 655.5 | ± | 3 640.1 | 2 550.0 | ± | 766.7 | 217 595.1 | ± | 14 508.7 |
| | | RSD | | 23% | RSD | | 30% | RSD | | 7% |
| | | - | / | 0.14192 | * | / | 0.01595 | *** | / | 0.00000 |
| **5 DAP** | **21 CT** | 7 435.3 | ± | 802.1 | 270.6 | ± | 15.3 | 17 592.6 | ± | 1 011.9 |
| | | RSD | | 11% | RSD | | 6% | RSD | | 6% |
| | **34 HT** | 6 755.5 | ± | 961.2 | 260.6 | ± | 53.3 | 28 585.4 | ± | 2 034.0 |
| | | RSD | | 14% | RSD | | 20% | RSD | | 7% |
| | | - | / | 0.30912 | - | / | 0.75714 | *** | / | 0.00005 |
| **26 DAP** | **21 CT** | 14 813.1 | ± | 2 759.9 | *NQ* | | | 33 648.6 | ± | 2 111.6 |
| | | RSD | | 19% | | | | RSD | | 6% |
| | **34 HT** | 7 990.2 | ± | 992.5 | *NQ* | | | 12 648.1 | ± | 984.5 |
| | | RSD | | 12% | | | | RSD | | 8% |
| | | ** | / | 0.00164 | | | | *** | / | 0.00000 |

| **sample type** | **temp** | average | | S.D. | av. | | S.D. | av. | | S.D. |
|---|---|---|---|---|---|---|---|---|---|---|
| | | **IAA** | | | **oxIAA** | | | **IAAsp** | | |
| **Pistil** | **21 CT** | 429.7 | ± | 171.1 | 292.9 | ± | 43.4 | 6.6 | ± | 0.8 |
| | | RSD | | 40% | RSD | | 15% | RSD | | 12% |
| | **34 HT** | 311.0 | ± | 11.2 | 298.9 | ± | 18.4 | 4.5 | ± | 1.3 |
| | | RSD | | 4% | RSD | | 6% | RSD | | 29% |
| | | - | / | 0.20350 | - | / | 0.81269 | - | / | 0.05361 |
| **5 DAP** | **21 CT** | 2 746.6 | ± | 108.8 | 11 409.7 | ± | 603.0 | 43.8 | ± | 2.4 |
| | | RSD | | 4% | RSD | | 5% | RSD | | 6% |
| | **34 HT** | 2 132.2 | ± | 83.1 | 15 045.6 | ± | 914.7 | 23.1 | ± | 2.1 |
| | | RSD | | 4% | RSD | | 6% | RSD | | 9% |
| | | *** | / | 0.00006 | *** | / | 0.00052 | *** | / | 0.00001 |
| **26 DAP** | **21 CT** | 14 152.9 | ± | 371.3 | 110 339.3 | ± | 6 979.5 | 87.5 | ± | 6.1 |
| | | RSD | | 3% | RSD | | 6% | RSD | | 7% |
| | **34 HT** | 3 904.8 | ± | 585.7 | 44 577.9 | ± | 3 869.0 | 2 087.7 | ± | 172.8 |
| | | RSD | | 15% | RSD | | 9% | RSD | | 8% |
| | | *** | / | 0.00000 | *** | / | 0.00000 | *** | / | 0.00000 |

| sample | | average | | S.D. | av. | | S.D. | av. | | S.D. |
|---|---|---|---|---|---|---|---|---|---|---|
| **type** | **temp** | **IAGlu** | | | **IAA-Glc** | | | **oxIAA-Glc** | | |
| **Pistil** | **21 CT** | *<LOD* | | | 143.3 | ± | 29.7 | 2 923.0 | ± | 169.6 |
| | | | | | RSD | | 21% | RSD | | 6% |
| | **34 HT** | *<LOD* | | | 150.8 | ± | 18.2 | 1 690.0 | ± | 83.5 |
| | | | | | RSD | | 12% | RSD | | 5% |
| | | | | | - | / | 0.69455 | *** | / | 0.00000 |
| **5 DAP** | **21 CT** | 62.9 | ± | 6.4 | 876.5 | ± | 57.3 | 43 524.6 | ± | 2 863.5 |
| | | RSD | | 10% | RSD | | 7% | RSD | | 7% |
| | **34 HT** | 54.4 | ± | 9.0 | 478.4 | ± | 68.5 | 43 785.4 | ± | 1 864.8 |
| | | RSD | | 17% | RSD | | 14% | RSD | | 4% |
| | | - | / | 0.20301 | *** | / | 0.00008 | - | / | 0.88867 |
| **26 DAP** | **21 CT** | 17.2 | ± | 1.8 | *NQ* | | | 788 378.5 | ± | 46 734.7 |
| | | RSD | | 11% | | | | RSD | | 6% |
| | **34 HT** | 96.1 | ± | 5.8 | *NQ* | | | 150 035.7 | ± | 11 672.8 |
| | | RSD | | 6% | | | | RSD | | 8% |
| | | *** | / | 0.00000 | | | | *** | / | 0.00000 |

NQ: not quantified; <LOD: below the limit of detection; CT: control temperature; HT: hight temperature; av.: average; S.D.; standard deviation; RSD: relative standard deviation (ratio of the standard deviation over the average). Asterisks indicate statistically significant difference in HT in a paired Student's t-test (t-test; *, **, and *** correspond to P-values of $0.05 > p > 0.01$, $0.01 > p > 0.001$, and $p < 0.001$, respectively). All measurements are as pmol/g FW.

*Supplementary Table S6.* ABA measurements (source data of Fig. 4)

| **sample** | | average | | S.D. |
|---|---|---|---|---|
| **type** | ***temp.*** | **ABA** | | |
| **Pistil** | **21 CT** | 1 101.6 | ± | 77.8 |
| | | RSD | | 7% |
| | **34 HT** | 513.6 | ± | 30.3 |
| | | RSD | | 6% |
| | | *** | / | 0.00000 |
| **5 DAP** | **21 CT** | 1 329.7 | ± | 32.0 |
| | | RSD | | 2% |
| | **34 HT** | 1 372.4 | ± | 63.8 |
| | | RSD | | 5% |
| | | - | / | 0.31725 |
| **26 DAP** | **21 CT** | 6 750.6 | ± | 283.6 |
| | | RSD | | 4% |
| | **34 HT** | 2 567.6 | ± | 100.3 |
| | | RSD | | 4% |
| | | *** | / | 0.00000 |

NQ: not quantified; CT: control temperature; HT: hight temperature; S.D.; standard deviation; RSD: relative standard deviation (ratio of the standard deviation over the average). Asterisks indicate statistically significant difference in HT in a paired Student's t-test (t-test; *** correspond to P-values of $p < 0.001$).

*Fig. S1*. Setup of growth temperatures in greenhouse chambers.

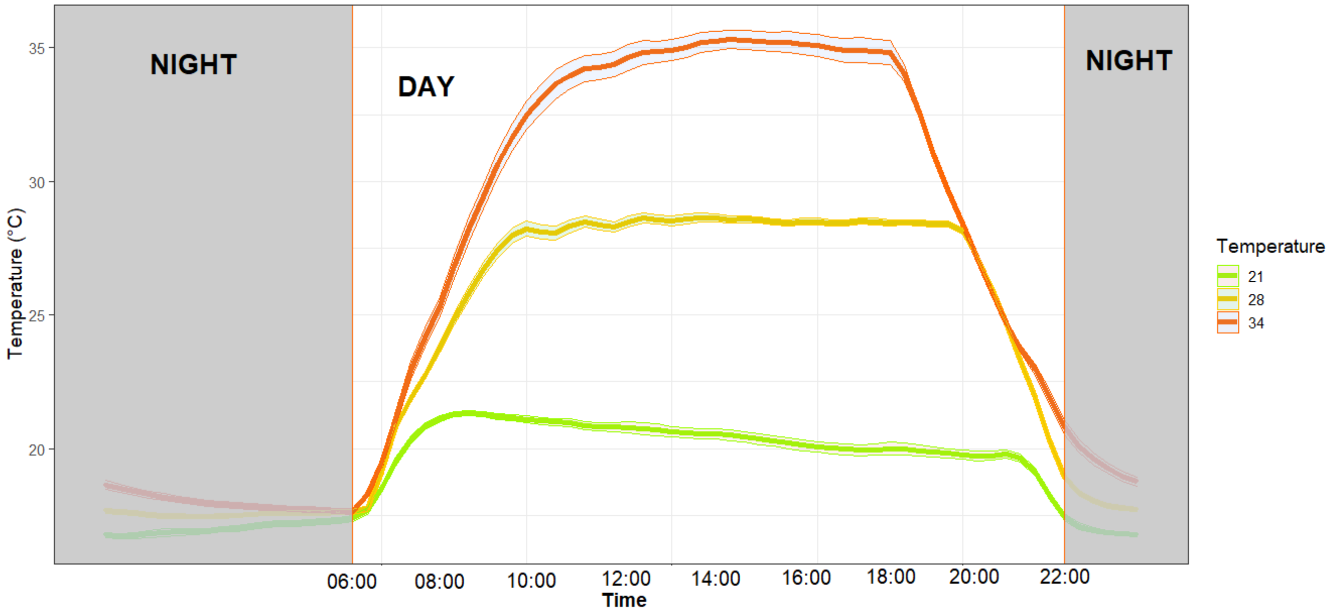


***Supplementary Figure S1*. Setup of growth temperatures in greenhouse chambers**

Three conditions were selected for the temperature stresses and control conditions. Temperature condition for control (CT, 21, green), mid (MT, 28, orange) and high temperature (HT, 34, red) chambers were set to 21 °C, 28 °C and 34 °C, respectively, with ramping of the temperature up and down by 4 °C per hour from night temperatures set at 18 °C for all conditions. Day period was set between 6:00 and 22:00 (light grey). The graph shows the mean temperatures (bold line) ± 95 % confidence interval in each greenhouse chamber measured throughout the experiment.

*Fig. S2*. High temperatures affect growth parameters of Brassica flowering plants

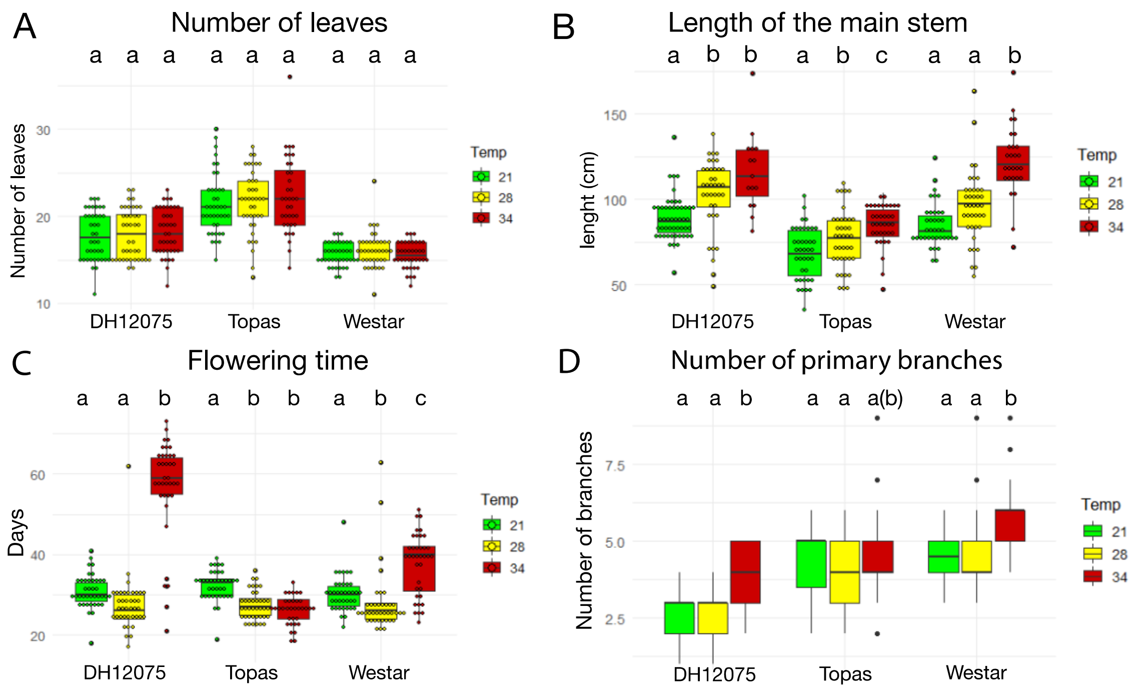


***Supplementary Figure S2*. High temperatures affect growth parameters of Brassica flowering plants**

Graphs

The number of leaves (A), length of the main stem (LMS, cm) (B), flowering time (FT, days) (C), and the number of primary branches (NB) (D) were quantified in DH12075, Topas and Westar cultivars at CT (21/18 ºC, green), MT (28/18 ºC, yellow) and HT (34/18 ºC, red).

Growth parameters are presented as boxplots (the box represents the interquartile range and the line inside the box represents the median). Each dot is an observation (A-C), or only outliers (D). The Pearson correlation coefficient between LSM, FT and NF, is shown in Supplementary Table S2. Boxes with the same letters (a, b, c) within each cultivar do no differ significantly ($p < 0.05$).

*Fig. S3.* The number of ovules is unchanged, but the development of the embryo is accelerated when plants are grown at high temperatures.

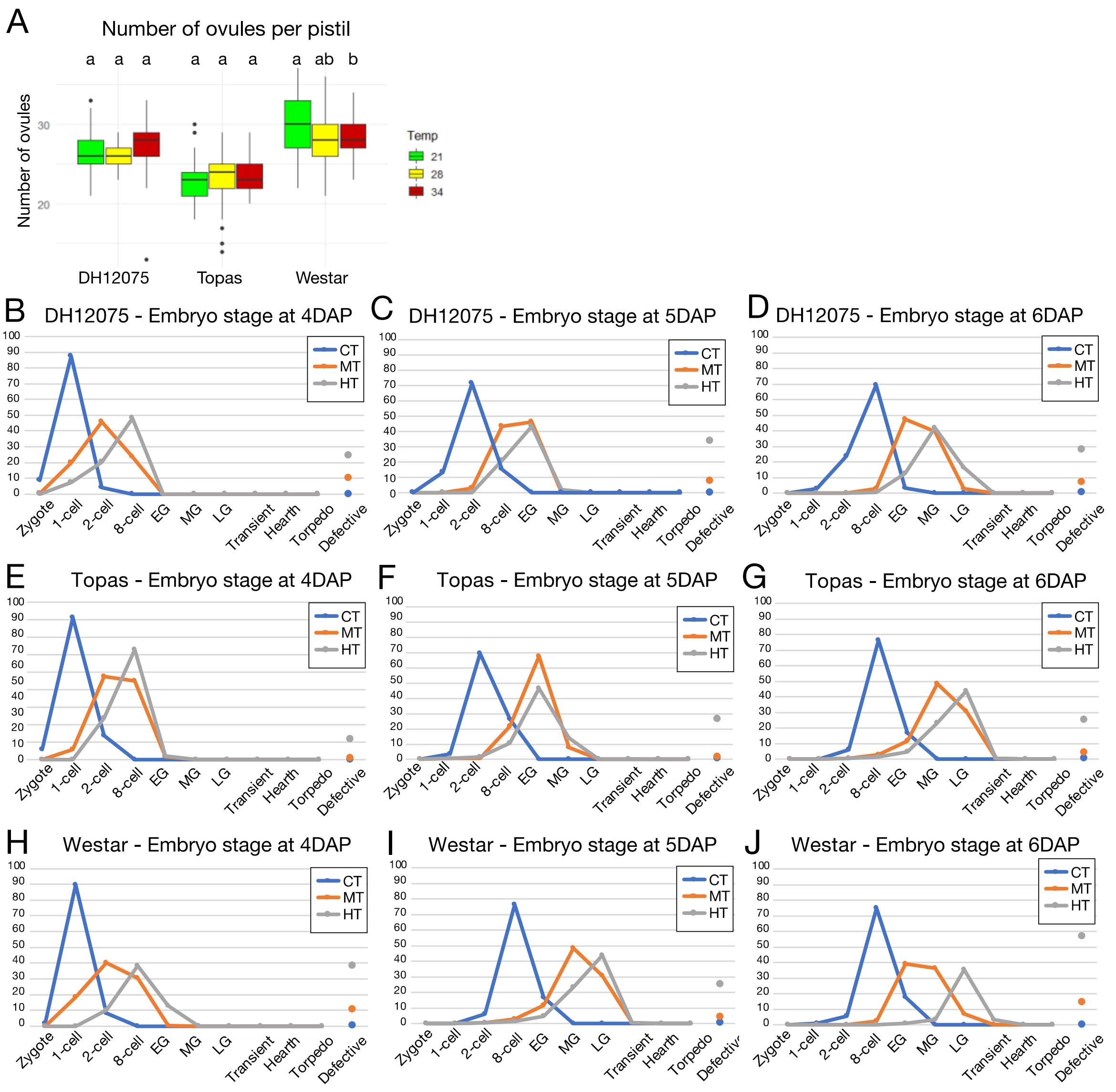


**_Supplementary Figure S3_. The number of ovules is unchanged, but the development of the embryo is accelerated when the plants are grown at high temperatures.**

(A) Graph displaying the number of ovules per pistil in DH12075, Topas and Westar cultivars at CT (21/18 °C, green), MT (28/18 °C, yellow) and HT (34/18 °C, red). Relates to Table 1. The number of ovules per pistil is presented as boxplots (the box represents the interquartile range and the line inside the box represents the median). Each dot indicates outliers. Boxes with the same letters (a, b, c) within each cultivar do no differ significantly ($p < 0.05$).

(B-J) Graphs displaying the distribution (as percentage) of embryonic development stages per silique at 4 DAP (B, E, H), 5 DAP (C, F, I) and 6 DAP (D, G, J) in DH12075 (B-D), Topas (E-G), and Westar plants (H-J) grown at CT (21/18 °C, blue), MT (28/18 °C, orange) and HT (34/18 °C, grey). EG, early globular; MG, mid-globular; LG, late globular embryos. Relates to Table 2.

*Fig. S4*. Pollen grain development is not affected by our stress growth conditions.

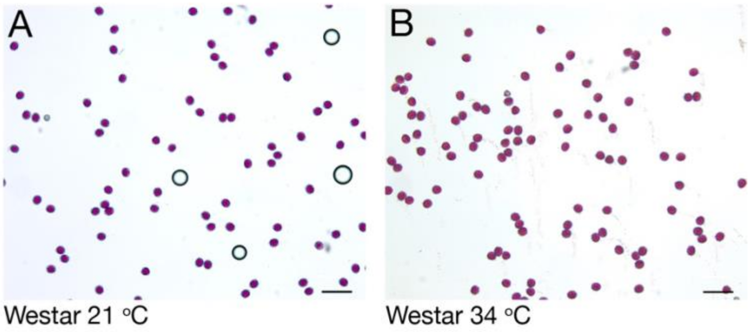


***Supplementary Figure S4*. Pollen grain development is not affected by our growth conditions.**

Pollen grains of Westar were assayed for viability with Alexander staining. No differences were observed in pollen viability from plants grown at 21 ºC (A) and 34 ºC (B). Scale bars represent 100 μm.

*Fig. S5*. High temperatures affect embryo development.

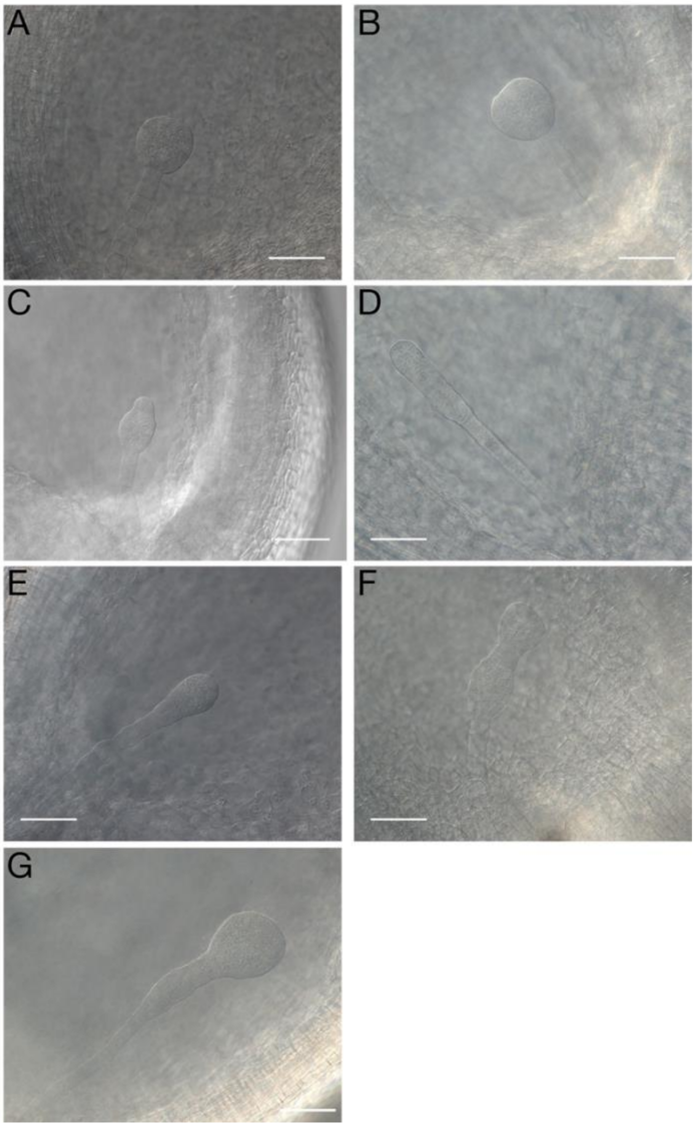


***Supplementary Figure S5*. High temperatures affect embryo development. Original pictures for the pictures presented in Fig 2.**

(A-B) Embryos at 6 DAP (A) and 7 DAP (B) from plants grown at CT (21/18 $^{o}$C).

(C-G) Range of defective embryos observed in *B. napus* plants grown at MT (28/18 $^{o}$C) and HT (34/18 $^{o}$C) between 6 and 8 DAP. Scale bars represent 100 $\mu$m.

*Fig. S6.* Silique growth rate is reduced at higher temperatures.

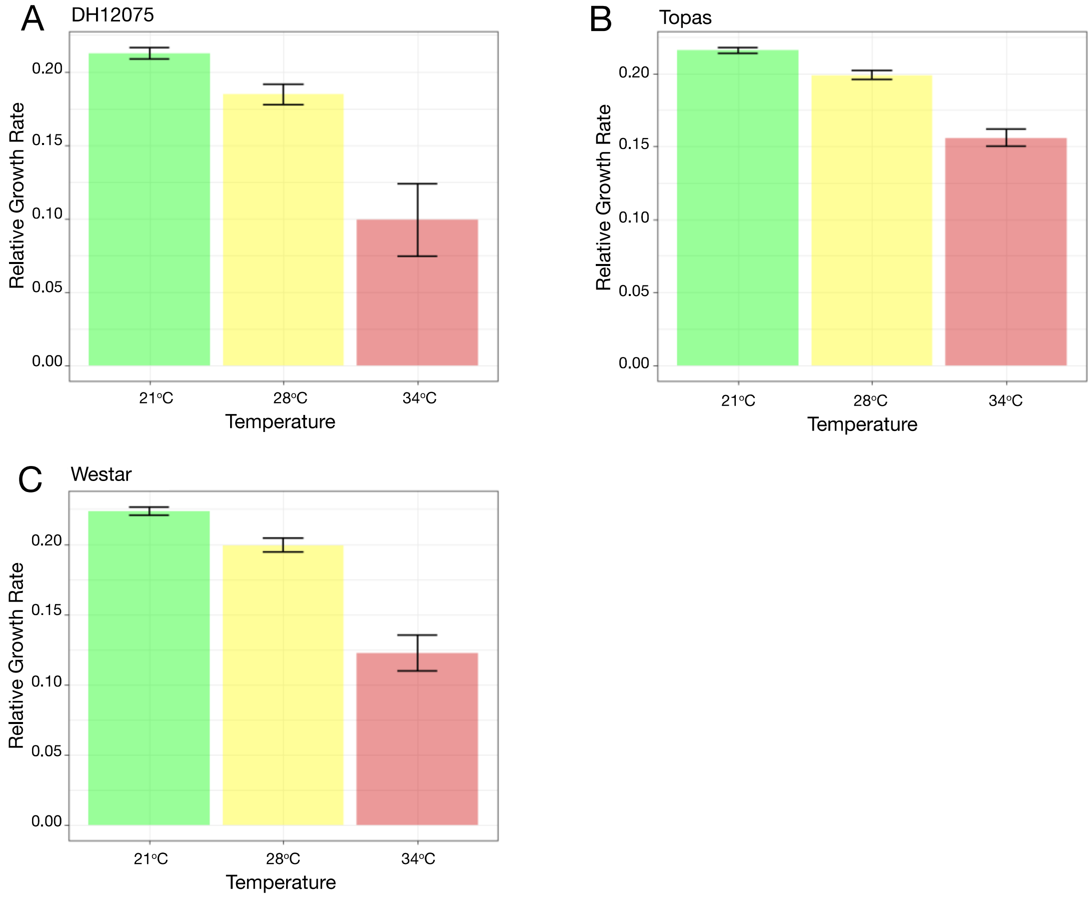


**Supplementary Figure S6. Growth rate of siliques are reduced by growth at higher temperatures.**
Graphs displaying the relative growth rate of siliques between 0 DAP to 11 DAP in DH12075 (A), Topas (B) and Westar (C) plants grown at CT (21/18 ºC, green), MT (28/18 ºC, yellow) and HT (34/18 ºC, red). Shown are barplots with 95 % confidence intervals.
